# Pre-neoplastic stromal cells drive BRCA1-mediated breast tumorigenesis

**DOI:** 10.1101/2021.10.20.465221

**Authors:** Kevin Nee, Dennis Ma, Quy H. Nguyen, Nicholas Pervolarakis, Jacob Insua-Rodríguez, Maren Pein, Yanwen Gong, Grace Hernandez, Hamad Alshetaiwi, Justice Williams, Maha Rauf, Kushal Rajiv Dave, Keerti Boyapati, Christian Calderon, Anush Markaryan, Robert Edwards, Erin Lin, Ritesh Parajuli, Peijie Zhou, Qing Nie, Sundus Shalabi, Mark A. LaBarge, Kai Kessenbrock

## Abstract

Women with germline mutations in *BRCA1* (BRCA1^+/mut^) have increased risk for developing hereditary breast cancer^1, 2^. Cancer initiation in BRCA1^+/mut^ is associated with pre-malignant changes in the breast epithelium including altered differentiation^3–5^, proliferative stress^6^ and genomic instability^7^. However, the role of the epithelium-associated stromal niche during BRCA1-driven tumor initiation remains unclear. Here, we show that the pre-malignant stromal niche promotes epithelial proliferation and BRCA1- driven cancer initiation in *trans*. Using single-cell RNAseq (scRNAseq) analysis of human pre-neoplastic BRCA1^+/mut^ and control breast tissues, we show that stromal cells provide numerous pro-proliferative paracrine signals inducing epithelial proliferation. We identify a subpopulation of pre-cancer associated fibroblasts (pre-CAFs) that produces copious amounts of pro-tumorigenic factors including matrix metalloproteinase 3 (MMP3)^8, 9^, and promotes BRCA1-driven tumorigenesis *in vivo*. Our gene-signature analysis and mathematical modeling of epithelial differentiation reveals that stromal-induced proliferation leads to the accumulation of luminal progenitor cells with altered differentiation, and thus contributes to increased breast cancer risk in BRCA1^+/mut^. Our results demonstrate how alterations in cell-cell communication can induce imbalances in epithelial homeostasis ultimately leading to cancer initiation. We anticipate our results to form the foundation for novel disease monitoring and therapeutic strategies to improve patient management in hereditary breast cancer. For example, pre-CAF specific proteins may serve as biomarkers for pre-cancerous disease initiation to inform whether radical bilateral mastectomy is needed. In addition, MMP inhibitors could be re-indicated for primary cancer prevention treatment in women with high-risk BRCA1 mutations.

## Main text

Stromal cells in the human breast are localized in tight association with epithelial ducts and lobules, where they regulate epithelial homeostasis through secretion of growth factors and extracellular matrix (ECM) molecules^10^. Here, we hypothesized that germline BRCA1^+/mut^ carriers exhibit alterations in the breast stromal niche, which promotes pre- malignant epithelial changes and cancer initiation. To define the heterogeneous stromal cell types and their communication with epithelium, we analyzed a cohort of non-tumorigenic breast mastectomies from BRCA1 germline mutation carriers (BRCA1^+/mut^; n=20) and non-mutation carriers (control; n=33) using a combination of scRNAseq, *in situ* analysis and functional *in vitro* and *in vivo* experiments. For scRNAseq (BRCA1^+/mut^: n=11; control: n=11), we first used differential centrifugation to enrich for breast epithelium^11^, then isolated epithelial (Lin-/EpCAM+) and stromal (Lin-/EpCAM-) cells by fluorescence activated cell sorting (FACS), and sequenced altogether 230,100 cells (**Fig. 1a, Extended Data Fig. 1a, Supplementary Table 1**). We used Seurat^12^ to identify the main cell types and their marker genes in a combined analysis of all samples (**Fig. 1b,c, Supplementary Table 2-3**). Importantly, cell type clusters contained cells from all individuals, and all samples demonstrated comparable quality metrics (**Extended Data Fig. 1b-e**). Within epithelium, we identified three cell types corresponding to basal (63,002 cells), luminal 1 (26,122 cells), and luminal 2 epithelial cells (28,045 cells), as previously described^13^. Within the epithelium-associated stroma, we found three main cell types corresponding to fibroblasts (55,428 cells; endothelial cells (31,819 cells), and pericytes^14^ (22,917 cells) (**Fig. 1b**).

**Fig. 1.**
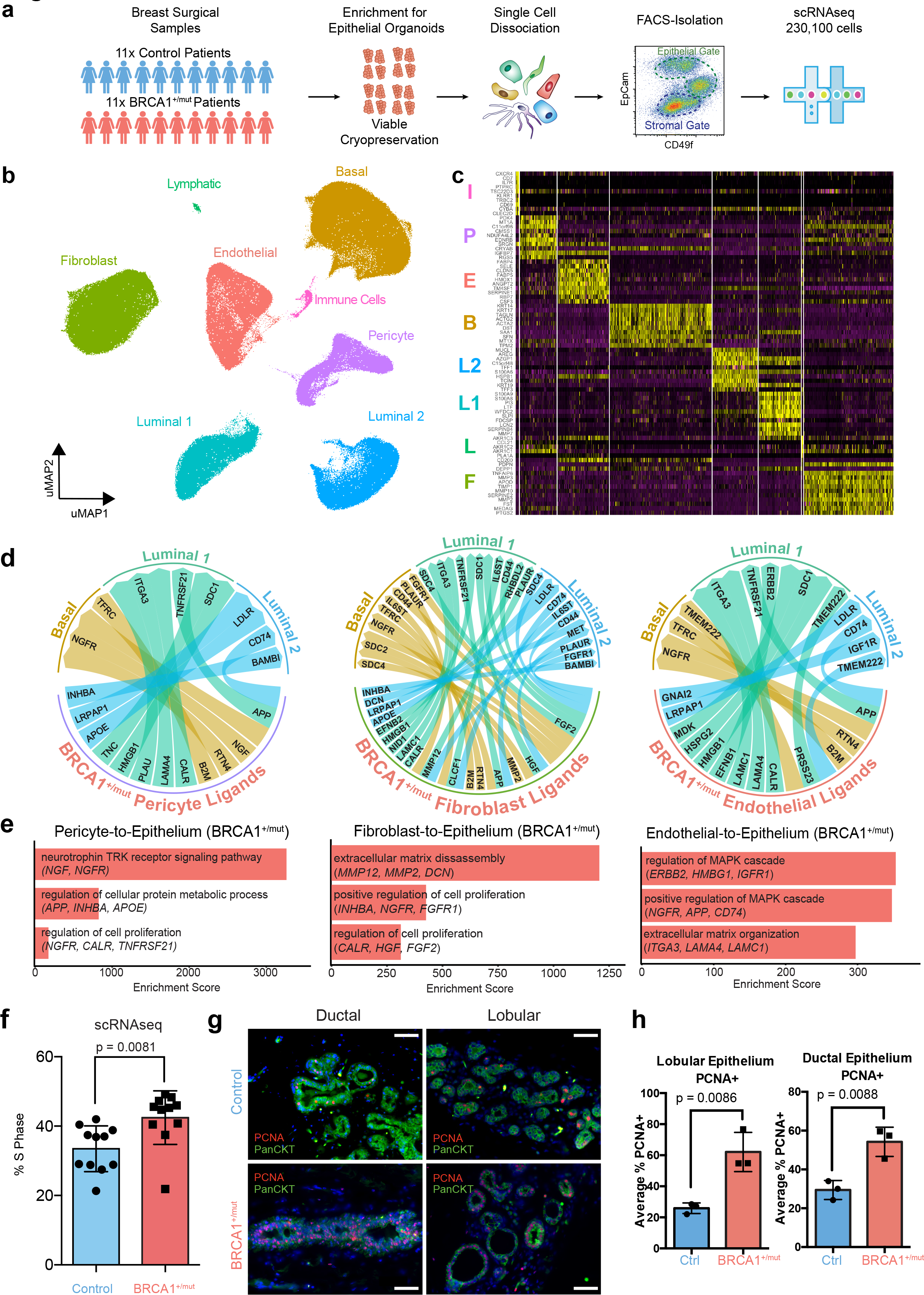
Single-cell transcriptomics analysis reveals overexpression of pro-proliferative stromal cues in the pre-neoplastic BRCA1^+/mut^ breast epithelial microenvironment. a) Schematic depiction of single cell analysis workflow using human breast tissue samples that are mechanically and enzymatically dissociated, isolated using flow cytometry to isolate stromal (EpCAM^-^) and epithelial (EpCAM^+^), which are separately subjected to scRNAseq. b) Integrated clustering analysis of n=11 control and n=11 BRCA1^+/mut^ scRNAseq dataset in uMAP projection showing the main cell types, i.e. the three epithelial cell types basal (B), luminal 1 (L1) and luminal 2 (L2), and the four stromal cell types fibroblast (F), pericytes (P), endothelial (E), lymphatic (L) as well as a cluster of immune cells (I). c) Top 10 marker gene heatmap for each cell type (rows=genes, columns=cells). The corresponding cell types are indicated with letter abbreviation. d) Circos plots showing ligand-receptor interactions enhanced in BRCA1^+/mut^ pericyte (left), fibroblast (center), and endothelial (right) ligands connecting with receptors of the three epithelial cell types basal, luminal 1 and luminal 2. e) Receptor-ligand interaction enrichment scores of GO Terms (GO-Biological Processes 2018) of BRCA1^+/mut^ pericyte (left), fibroblast (center), and endothelial (right) ligands, and epithelial receptors are shown. f) Bar chart shows the percentage of each patient’s (control: n=11; BRCA1^+/mut^: n=11) epithelial cells in S phase as identified by Seurat cell-cycle scoring analysis. Unpaired, two-tailed, t-test with equal SD calculates p-value = 0.0081. g) Representative images from IF analysis of pan-cytokeratin and PCNA immunostainings showing ductal and lobular regions in control and BRCA1^+/mut^ samples. Scale bar = 50um. h) Bar graphs showing average percentage of PCNA+ cells in n=5 regions each from control (n=3) and BRCA1^+/mut^ (n=3) patients by *in situ* IF analysis of lobular and ductal areas.

Since fibroblasts and pericytes have been historically difficult to distinguish^15–17^, we next defined molecular differences and commonalities between these breast stromal cell types through differential marker gene expression and gene ontology (GO) term analysis (**Extended Data Fig. 2a-e**). Among the commonalities were genes associated with mesenchymal biology including *EDNRB, PDGFRB, ZEB2* and *COL4A1*^18^. Key differences were ECM molecules (*COL1A2*) and proteolytic remodelers (*MMP2*, *MMP3*, *MMP10*) in fibroblasts, and actin-binding (*TAGLN*, *ACTA2*) and endothelial factors (*PROCR*, *ESAM*, *MCAM, KCNE4*) in pericytes highlighting their vasocontractile function (**Extended Data Fig. 2b**). Importantly, we found that cell surface markers *PROCR* and *PDPN* differentially labelled pericytes and fibroblasts (**Extended Data Fig. 2f**), which allowed us to develop a FACS strategy to specifically enrich for pericytes (PROCR+PDPN-) and fibroblasts (PDPN+PROCR^mid^) (**Extended Data Fig. 2g-i**). Our approach for selective isolation of fibroblasts and pericytes from human tissues allows for prospective functional analyses and may improve therapeutic approaches utilizing the regenerative capacity of pericytes^15^.

**Fig. 2.**
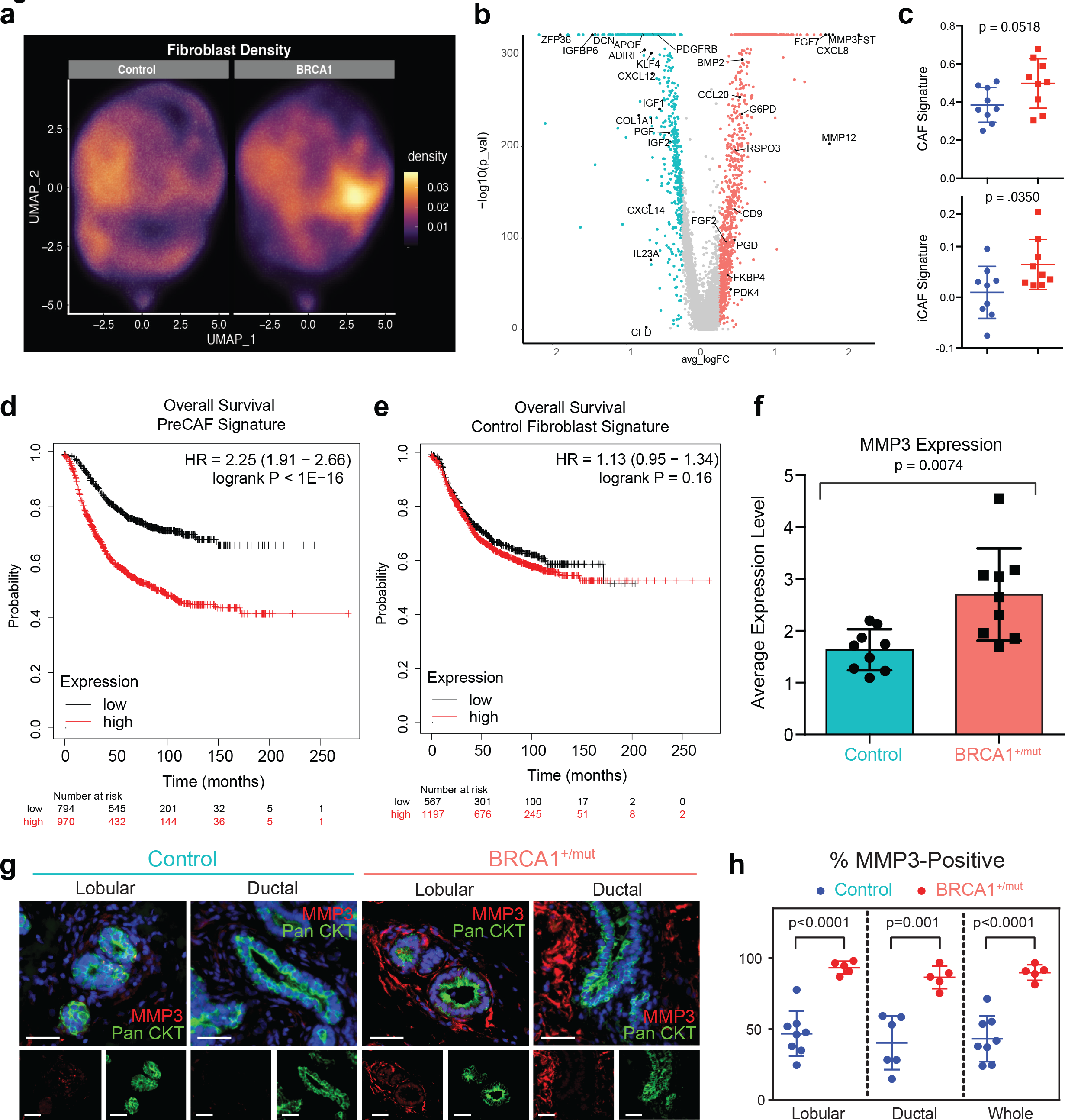
Expansion of CAF-like, MMP3-expressing fibroblasts in pre-malignant BRCA1^+/mut^ breast tissues. a) uMAP projection of cell density by control and BRCA1^+/mut^ fibroblasts. b) Volcano plot with all differentially expressed genes between control and BRCA1^+/mut^ fibroblasts. Top 50 BRCA1^+/mut^ genes were used to define pre-CAF signature. Top 50 control genes define our control Fibroblast Signature. c) Gene signature scoring of all control and BRCA1^+/mut^ fibroblasts for cancer associated fibroblast (CAF) and iCAF signatures. P-value determined by welch two sample t-test. d) Overall survival Kaplan-Meier analysis of n=1764 patients with a high (n=970) and low (n=794) expression of the pre-CAF signature. e) Overall survival Kaplan-Meier analysis of n=1764 patients with a high (n=1197) and low (n=567) expression of the control fibroblast signature. f) Boxplot of MMP3 expression in control and BRCA1^+/mut^ fibroblasts. *P* values are determined using a Mann-Whitney test. g) In situ immunofluorescence (IF) analysis of lobular and ductal regions of breast epithelium in control and BRCA1^+/mut^ human tissue sections. Scale bar = 50 µm. h) Dot plot showing percentage of MMP3-positive stromal cells in control (blue) and BRCA1^+/mut^ (red) samples as quantified from IF stainings. *P*-values were determined using unpaired t-tests.

Since stromal cells play key roles in regulating epithelial homeostasis through paracrine and juxtacrine interactions^10^, we next explored ligand-receptor interactions that displayed enhanced expression patterns in BRCA1^+/mut^ compared to control samples (**Supplementary Table 4**). Reassuringly, fibroblasts and pericytes were found to produce several collagen/integrin interactions with epithelium, which were underrepresented in BRCA^+/mut^ compared to control (**Extended Data Fig. 3**). Intriguingly, we found several tumor-promoting growth factors enriched in BRCA1^+/mut^, including *FGF2*^19^ and *HGF*^20^ from fibroblasts, and *NGF*^21^ and *INHBA*^22^ from pericytes (**Fig. 1d**). GO term analysis showed that BRCA1^+/mut^ samples exhibit an overall increase in pro-proliferative cues from both pericytes and fibroblasts, while endothelial cells exhibited increases in inducing

**Fig 3.**
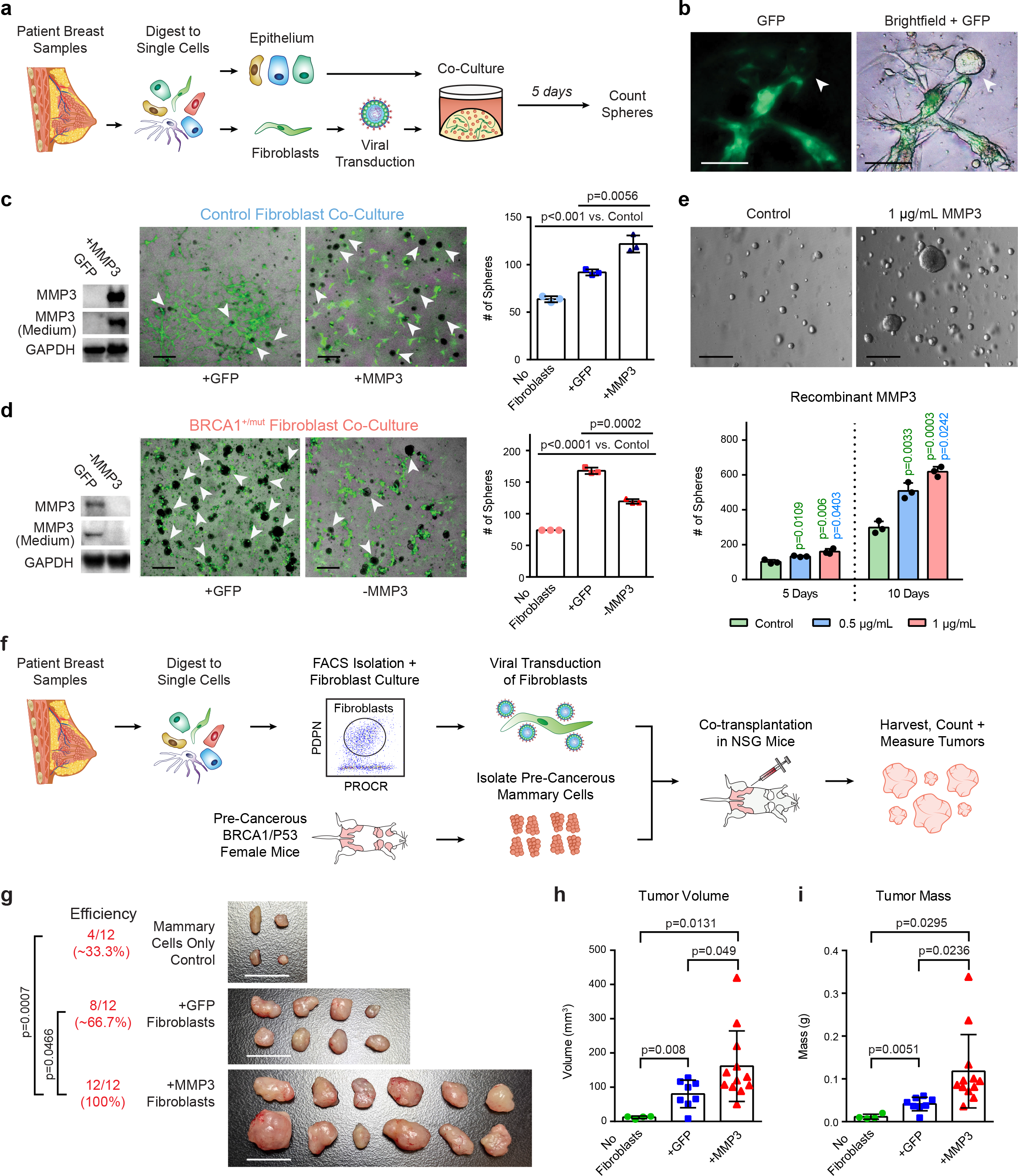
MMP3-expressing pre-CAFs promote breast epithelial growth *in vitro* and BRCA1-mediated breast tumorigenesis *in vivo*. **a)** Schematic primary human 3D co-culture using FACS-isolated epithelial cells and fibroblasts modulated using lentiviral transduction. **b)** Representative images depicting green fluorescent protein (GFP) expression in transduced fibroblasts in close proximity with epithelial organoids (arrow). Scale bar = 100 μm. **c)** Co-cultures (5 days) of 4,000 breast epithelial cells seeded alone (no fibroblasts) or with 1x10^4^ control fibroblasts transduced with lentivirus (+GFP) or transduced to express MMP3 and GFP (+MMP3). Western blots show increased expression of MMP3 in cells and cultured supernatant of +MMP3 fibroblasts (left). Representative merged bright field and GFP images of co-cultures (scale bar= 400 μm) with arrows indicating mammospheres (GFP-negative). Bar charts (right) represent number of mammospheres quantified per well; values expressed as mean ± SD from three separate experiments with three triplicate wells from each. *P*-values were determined using unpaired t-tests. **d)** Co-cultures (5 days) of 4,000 breast epithelial cells seeded alone (no fibroblast) or with 1x10^4^ BRCA1^+/mut^ fibroblasts transduced with lentivirus to express CRISPR- Cas9 and MMP3 gRNA (-MMP3) or GFP only (+GFP) vectors. Western blots show decreased expression of MMP3 in cells and medium of MMP3 knock out fibroblast cultures (left). Representative overlay bright field and GFP images of co-cultures (scale bar= 400 μm) with arrows indicating mammospheres (GFP-negative). Bar charts (right) represent number of mammospheres quantified in each well; values expressed as mean ± SD from triplicates of three independent experiments. *P*- values were determined using unpaired t-tests. **e)** 1x10^4^ FACS-isolated epithelial cells from patient sample “Control 36” were seeded in Matrigel and treated with 0.5 µg/mL or 1 μg/mL recombinant MMP3. Number of mammospheres was quantified after 5 and 10 days. Representative bright field images of mammospheres after 10 days of culture (scale bar= 400 μm). Bar chart values are expressed as mean ± SD from triplicates of three independent experiments. *P*-values were determined using unpaired t-tests. **f)** Schematic of mouse model for evaluating effects of stromal fibroblasts (Control 27) on BRCA1-mediated breast tumorigenesis *in vivo*. **g)** Images of dissected tumors after six weeks of growth with reported tumor formation efficiencies. Scale bar= 1 cm. *P*-values were determined using a one- sided Fisher’s Exact test. **h)** Volumes of dissected tumors. Values are represented as mean ± SD. control n= 4; +GFP n=8, MMP3 n=12. *P*-values were determined using unpaired t-tests. **i)** Masses of dissected tumors. Values are represented as mean ± SD. control n= 4; +GFP n=8, +MMP3 n=12. *P*-values were determined using unpaired t-tests.

MAPK signaling (**Fig. 1e**), suggesting that alterations in the stromal niche drive epithelial proliferation in pre-malignant breast tissues. Indeed, our cell-cycle scoring analysis^23^ identified an increased percentage of BRCA1^+/mut^ epithelial cells in S phase (**Fig. 1f**). To validate this *in situ*, we performed immunofluorescence staining for PCNA in additional control (n=3) and BRCA1^+/mut^ (n=3) samples, which confirmed a ∼2-fold increased number of PCNA-positive epithelial cells (**Fig. 1g-h**).

Our ligand-receptor analysis revealed NGF as a pro-proliferative factor in BRCA^+/mut^ pericytes interacting with NGFR on basal cells (**Fig. 1d**). We performed a more detailed clustering analysis of vascular (endothelial and pericyte) cell states (**Extended Data Fig. 4a,b),** which confirmed that NGF was significantly overexpressed in BRCA^+/mut^ pericytes **(Extended Data Fig. 4c, Supplementary Table 5).** NGF is known to induce proliferation in cancer cells^21^, however it is not known to act as a microenvironmental growth factor in the pre-cancerous breast. This interaction was supported by our FACS results that only basal cells express NGFR (**Extended Data Fig. 4d**). To functionally test this, we investigated whether FACS-isolated NGFR+ basal cells display increased proliferation when cultured with exogenous NGF in mammosphere formation assays^9^. Indeed, addition of NGF induced significantly increased number and size of mammospheres of basal, but not luminal cells (**Extended Data Fig. 4e-f**), and enhanced mammary branching morphogenesis branching morphogenesis^24^ in a physiologically relevant ECM hydrogel assay^25^ (**Extended Data Fig. 4h,i)**. Together, these findings revealed previously unrealized molecular mechanisms underlying the microenvironmental induction of epithelial proliferation in pre-neoplastic BRCA1^+/mut^ breast tissues.

**Fig. 4.**
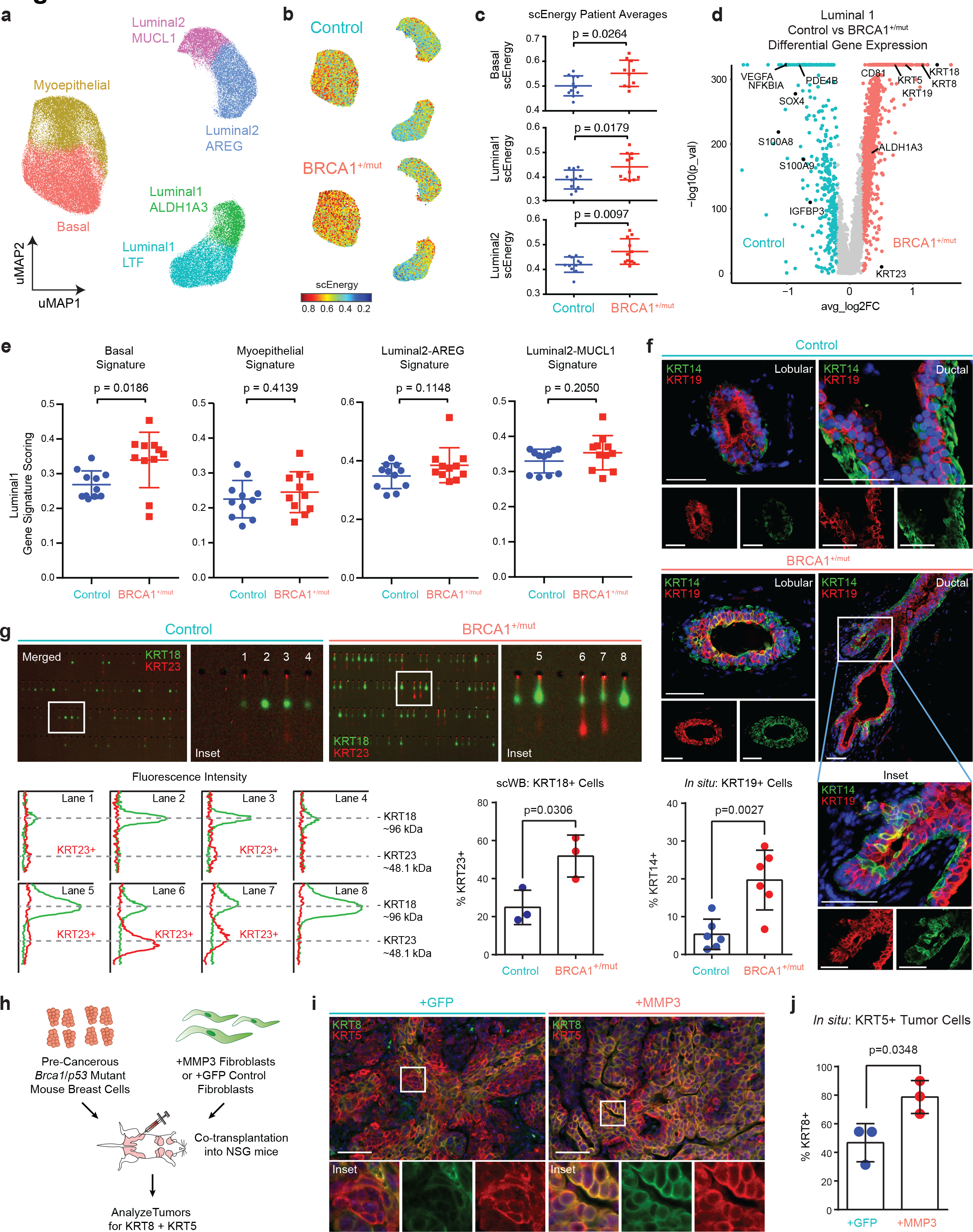
Analysis of epithelial differentiation reveals accumulation of a luminal progenitor subset with altered differentiation in BRCA1^+/mut^. a) Unbiased clustering using uMAP projection of all patient epithelial cells. Cells are labeled by mammary epithelial cell state classification as indicated. b) uMAP feature plots displaying single cell energy in facetted plots for control (upper plot) and BRCA1^+/mut^ cells (lower plot). c) scEnergy distributions across basal, luminal 1, and luminal2 cell types from control and BRCA1^+/mut^ samples. Mean scEnergy values from individual patients are plotted. P-value determined by welch two sample t-test. d) Volcano plot displaying genes differentially expressed between control and BRCA1^+/mut^ luminal 1 epithelial cells. e) Gene signature scoring of luminal 1 cells from control and BRCA1^+/mut^ epithelial cells for basal, myoepithelial, luminal 2-AREG, and luminal 2-MUCL1 marker gene signatures. P-value determined by Welch two sample t-test. f) *In situ* immunofluroescence analysis of KRT14/KRT19-double positive cells of lobular and ductal regions in control and BRCA1^+/mut^ tissues with representative images shown. Bar chart (bottom left) indicates the percentage of K14/K19-double positive cells per field of view in control (n=6) and BRCA1^+/mut^ (n=6). Values are expressed as mean ± SD quantified from at least 5 random fields per patient sample. *P*-value was determined using an unpaired t-test. g) Single cell Western blot (scWB)-based quantification of the percentage of KRT23+ in FACS-isolated luminal epithelial cells from n=3 control, and n=3 BRCA1^+/mut^. Images are representative regions of scWB chips post electrophoresis and antibody probing. Bar chart values are represented as mean ± SD from at least 1000 cells/individual; n=3 control, and n=3 BRCA1^+/mut^. h) Schematic workflow for analyzing co-transplant tumors from *in vivo* study described in Fig. 3f**-i** for KRT8 and KRT5 expression. i) Representative immunofluorescence images of sections from tumors produced from +GFP control or +MMP3 fibroblast transplants. Yellow staining indicates tumor cells double positive for KRT8 and KRT5. Scale bar= 50 µm. j) Bar chart is a quantification of the percent of KRT5 cells positive from KRT8. Values are represented as mean ± SD from counts from three different tumors per group with at least five random fields per tumor tissue. *P*-value was determined by an unpaired t-test.

Fibroblasts are critical niche cells that regulate normal breast epithelial homeostasis through secretion of growth factors and ECM molecules^10^ and contribute to tumor progression as CAFs^26^. Our subset fibroblast cell density analysis showed distinct changes in transcriptional space between control and BRCA1^+/mut^ (**Fig. 2a**). We next performed differential gene expression analysis revealing striking differences in gene expression between BRCA1 and control fibroblasts (**Fig. 2b, Supplementary Table 6**) Interestingly, further gene scoring analysis showed elevated expression of CAF^27^ and iCAF^28^ signature genes (**Fig. 2c**) in BRCA1^+/mut^ fibroblasts, suggesting that a subset of BRCA1^+/mut^ fibroblasts consistently assume a “pre-CAF” phenotype at the pre-malignant stage (**Extended Data Fig. 5a**). Kaplan-Meier survival analysis^29^ of a cohort of 1,809 patients using the top 100 marker genes showed strong association of the pre-CAF, but not the control gene signature with poor outcome (**Fig. 2d,e**) underlining the potential disease relevance of this pre-CAF population.

In particular, the secreted protease matrix metalloproteinase 3 (MMP3) was one of the top pre-CAF markers found to be elevated in BRCA1^+/mut^ fibroblasts compared to control across all individuals (**Fig. 2f**). We and others previously demonstrated that MMP3 can promote breast cancer initiation through Wnt signaling^8, 9^, reactive oxygen production and increased genomic instability^30^, as well as during aging^31^, however the role of MMP3 in human BRCA1^+/mut^ pathology remains unclear. To validate whether MMP3 expression is increased in BRCA1^+/mut^ fibroblasts at the protein level *in situ*, we performed immunofluorescence analysis on additional control (n=8) and BRCA1^+/mut^ (n=5) breast tissue samples. This revealed an expansion of MMP3-positive stromal cells in close proximity to epithelial structures in BRCA1^+/mut^ tissues (**Fig. 2g,h,** Extended Data **Fig. 5b**) suggesting a direct link of tumor-promoting MMP3 to increased breast cancer risk in human BRCA1^+/mut^.

To functionally determine the effects of fibroblast-derived MMP3 on human breast epithelial biology, we established a 3D stromal-epithelial co-culture assay using primary human breast fibroblasts and mammary epithelial cells (MECs) (**Fig. 3a,b**). We next used lentiviral MMP3 overexpression in control fibroblasts (+MMP3) which revealed significantly increased growth compared to control-GFP only fibroblasts (+GFP) (**Fig. 3c, Extended Data Fig. 6a-d**). Conversely, deleting MMP3 using CRISPR/Cas9-mediated knockout in BRCA1^+/mut^ fibroblasts (-MMP3) resulted in significant reduction of mammosphere growth (**Fig. 3d**). To determine whether MMP3 directly promotes epithelial growth, we next added recombinant MMP3 to the mammosphere growth assay. We found that exogenous MMP3 was sufficient to induce increased mammosphere growth in a concentration-dependent manner (**Fig. 3e, Extended Data Fig. 6e,f**).These results show that fibroblast-derived MMP3 acts in *trans* to promote breast epithelial growth in the human system.

To evaluate the effect of fibroblast-derived MMP3 on tumor initiation *in vivo*, we established a fibroblast-epithelial co-transplantation mouse model for BRCA1-driven tumor initiation (**Fig. 3f**). In brief, we first isolated pre-cancerous mammary cells from *Brca1^f11/f11^p53^f5&6/f5&6^Cre^c^* mice^5^. We also FACS-isolated (PDPN+) human breast fibroblasts and lentivirally modulated these to express GFP only (+GFP) or both MMP3 and GFP (+MMP3) (**Extended Data Fig. 6g**). We then performed orthotopic co-transplantation into immunocompromised mice in three experimental groups (n=12 each): 1. mammary cells only (control), 2. Mammary cells with control +GFP fibroblasts, 3. mammary cells with +MMP3 fibroblasts. After six weeks, we found strongly increased tumor initiation frequency in the +MMP3 group (12/12) compared to mammary cells only (4/12), and the +GFP control groups (8/12) demonstrating that fibroblast-derived MMP3 promotes BRCA1-mediated tumor initiation (Fig. 3g). Additionally, comparing tumor volume and mass showed significantly increased tumor growth in the +MMP3 group compared to both control groups (Fig. 3h**,i**). These results demonstrate that fibroblast-derived MMP3 drives BRCA1-mediated breast tumorigenesis in a paracrine fashion *in vivo*.

To determine whether stromal-induced proliferation alters epithelial differentiation in pre-malignant BRCA1^+/mut^ tissues, we next performed subset epithelial cell clustering and classified all epithelial cells on the cell state level as previously described^1^^3^ (Fig. 4a**, Extended Data** Fig. 7a**, Supplementary Table 8**). To identify cell states that have progenitor capacity we used a statistical physics-based approach that quantifies increased cell state transition probabilities as a single cell energy (scEnergy)^32^, and found that BRCA1^+/mut^ basal, luminal1, and luminal2, epithelial cells had significantly higher scEnergy than their control counterparts (Fig. 4b**-c**). In support of this notion, BRCA1^+/mut^ basal epithelial cells exhibited enhanced transcription of hallmark luminal cytokeratins (*e.g.*, KRT18, KRT8, KRT19) (**Extended Data** Fig. 7b), suggesting a phenomenon of altered differentiation. To validate this, we performed single cell western blot (scWB) analysis for dual expression of luminal (KRT19) and basal (KRT14) markers in isolated basal epithelial cells, which revealed increased percentage of KRT19/KRT14-double positive basal cells in BRCA1^+/mut^ (**Extended Data** Fig. 7c**-e)**.

Previous studies have implicated luminal progenitors (*i.e.*, luminal 1) as the cancer cell-of-origin in BRCA1^+/mut^- associated breast cancer^3, 4, 33–35^. In contrast to previous work, we did not detect an expansion in luminal 1 cells in BRCA1^+/mut^. However, our detailed analyses within luminal 1 cells revealed that basal hallmark genes (*e.g.,* KRT5, KRT14) as well as luminal progenitor genes (*e.g.,* ALDH1A3) were upregulated in BRCA1^+/mut^ luminal 1 cells (**Fig. 4d**). Furthermore, luminal 1 cells exhibited increased gene scores for basal, but not other epithelial cell type-associated gene signatures (**Fig. 4e**). BRCA1^+/mut^ Luminal1-ALDH1A3 epithelial cells were also found to have altered differentiation in BRCA1^+/mut^ patients, with increased expression of hallmark basal epithelial cytokeratins (e.g. *KRT5*) (**Fig. 4d**), and exhibited higher expression of basal cell state gene signature (**Fig. 4e; Supplementary Table 7)**. This observation was corroborated using *in situ* immunofluorescence as the percentage of KRT14/KRT19-double positive luminal cells was significantly increased in BRCA1^+/mut^ tissues (**Fig. 4f, Extended Data Fig. 8a,b, Supplementary Table 9)**. The same subset of luminal1-ALDH1A3+ was also found to express high levels of KRT23. To validate this finding, we performed single-cell Western blot (scWBs) analysis of epithelial cells isolated from control (n=3) and BRCA1^+/mut^ tissues (n=3), showing that BRCA1^+/mut^ patients have a greater percentage of luminal epithelial positive for KRT23 (**Fig. 4g)**. This was further corroborated by *in situ* immunofluorescence analysis showing increased numbers of KRT23-positive luminal cells in BRCA1^+/mut^ (**Extended Data Fig. 9a,b, Supplementary Table 10**).

To determine if stromal MMP3 expression directly induces altered differentiation, we performed *in situ* analysis on tumors derived from co-transplantation of MMP3- overexpressing or control (GFP) fibroblasts, which revealed a significant increase of KRT5+/KRT8+ double-positive tumor cells when stromal MMP3 is overexpressed (**Fig. 4h-j**). Together our data shows the expansion of luminal cells with altered basal-like differentiation, which is in line with previous observations^35, 36^. Importantly, our work revealed that the altered differentiation phenotype can be induced in trans by stromal cells, for example through secreted MMP3.

Finally, we sought to evaluate whether stromal cell-induced epithelial proliferation is likely the cause for progenitor cell expansion and increased breast cancer risk in BRCA1^+/mut^. BRCA1 haploinsufficiency is associated with increased genomic instability during proliferation^37^, thus increased proliferation may further accelerate the process of acquiring loss of BRCA1 heterozygosity or second oncogenic hits. To this end, we used a mathematical modeling approach simulating the population dynamics of cancer progenitors based on a previously developed mammary stem and progenitor hierarchal model^38^. We simulated the development of sequential mutations in BRCA1 and other oncogenes (e.g. p53) during cancer initiation (**Extended Data Fig. 10a**). Our results predict that 2-fold stromal-induced proliferation increase leads to marked accumulation of a potential cancer progenitor population (**Extended Data Fig. 10b-e**), which is in line with our finding of uncommitted luminal progenitor cell expansion in BRCA1^+/mut^ (**Fig. 4d-g**). To achieve a more realistic prediction of cancer risk over human lifespan, we used a random mutation model^39^ that assumes acquired mutations induce stochastic changes in cancer cell fitness^40^. Our model predicts that 2-fold increase in proliferation leads to markedly higher in overall risk of cancer (**Extended Data Fig. 10f**). This shows that stromal cell-induced proliferation is likely to be directly linked with progenitor cell expansion and increased breast cancer risk in BRCA1^+/mut^. Taken together, our work identifies pre-malignant alterations in stromal cell populations, which provide a conducive, pro-tumorigenic niche in human BRCA1^+/mut^ inducing the expansion of KRT23-positive subpopulation of luminal progenitors (**Extended Data Fig. 10d,e**).

Many other studies have characterized BRCA1^+/mut^ pre-neoplastic tissues including using scRNAseq^3, 4, 33, 36, 41, 42^. While these studies primarily focused on epithelial cells, our current work revealed the distinct pre-neoplastic changes within various stromal cell populations such as preCAFs. Furthermore, our work shows that stromal cells drive hereditary breast cancer in *trans*, which may pave the way towards novel disease monitoring and therapeutic strategies to improve of BRCA1^+/mut^ patient management. For example, our results indicate that MMPs, MMP3 in particular, may be a promising drug target for primary cancer prevention in BRCA1^+/mut^ carriers. Although MMP inhibitors have been tested as anti-cancer drugs in previous clinical trials with mostly disappointing results, poor study design focusing on late stage cancer patients may have contributed to the results of these trials^43^. Our study implies that targeting stromal-epithelial interactions, for example with MMP inhibitors, should be investigated for primary cancer prevention treatment in women with high risk BRCA1 mutations.

## MATERIALS AND METHODS

### Collection and processing of primary human breast tissues

Non-tumorigenic control and BRCA1^+/mut^ breast tissue samples were acquired from University of California, Irvine Chao Family Comprehensive Cancer Center under IRB protocol UCI 17-05, from the cooperative human tissue network (CHTN) and City of Hope Cancer Center under IRB protocol #17185 (see **Supplementary Table 1**). The main inclusion criteria for both control and BRCA1^+/mut^ samples was that they were histopathologically normal tissues (i.e., non-tumorigenic samples from reduction mammoplasty, prophylactic mastectomy, or contra-lateral mastectomy surgeries). For samples included in single-cell sequencing, the respective BRCA1 mutation or absence thereof was confirmed by DNA sequencing; for samples procured through CHTN, confirmation of BRCA1 mutations was provided by the respective clinical center. Tissue samples were processed as previously reported^13^. In brief, surgical specimens were washed in PBS, mechanically dissociated with scalpels, digested with 2 mg/mL collagenase I (Life Technologies, 17100-017) in DMEM (Corning, 10-013-CV) overnight. Next, the samples were treated with 20 U/mL DNase I (Sigma-Aldrich, D4263-5VL) for 5 min, and enriched for epithelium-rich tissues tissues and closely associated stroma by differential centrifugation using two pulses of 150 g. Supernatant was collected and centrifuged for 5 min x 500 g to isolate epithelial tissue chunks in the pellet. These were viably cryopreserved in DMEM with 50% FBS (Omega Scientific, FB-12) and 10% DMSO (v/v) before processing into single cells for scRNAseq or functional cell-based assays.

### Single-cell transcriptomics

Primary human organoids were digested with 0.05% trypsin (Corning, 25-052-CI) containing 20 U/mL DNase I to generate single cell suspensions. Cells were stained for FACS using fluorescently labeled antibodies for CD31 (eBiosciences, 48-0319-42), CD45 (eBiosciences, 48-9459-42), EpCAM (eBiosciences, 50-9326-42), CD49f (eBiosciences, 12-0495-82), SytoxBlue (Life Technologies, S34857). Only samples with at least 80% viability as assessed using SytoxBlue with FACS were included in this study. For scRNAseq, we excluded doublets, dead cells (SytoxBlue+), lin+ (CD31/CD45+), and isolated epithelial (EPCAM+) and stromal (EPCAM-) cells separately (complete list of antibodies in **Supplementary Table 11**). Flow cytometry sorted cells were washed with 0.04% BSA in PBS and suspended at approximately 1000 cells/µL. Each patient tissue sample was generated as an individual scRNAseq library. Generation of libraries for 10x Genomics v1 chemistry (sample IDs: control 1; BRCA1^+/mut^ 1) was performed following the Chromium Single Cell 3’ Reagents Kits User Guide: CG00026 Rev B. Library generation for 10x Genomics v2 chemistry (sample IDs: control 2-11; BRCA1+/mut 2-11) were performed following the Chromium Single Cell 3’ Reagents Kits v2 User Guide: CG00052 Rev B. cDNA library quantification was performed using Qubit dsDNA HS Assay Kit (Life Technologies, Q32851) and high-sensitivity DNA chips (Agilent. 5067- 4626). Quantification of library construction was performed using KAPA qPCR (Kapa Biosystems KK4824). The Illumina HiSeq4000 and NovaSeq6000 platform was used to achieve an average of 50,000 reads per cell. Alignment of sequencing results was performed using 10x Cell Ranger v3.1 to the GRCh38 reference.

### Seurat analysis of scRNAseq data

The Seurat pipeline (version 4.0.0) was used for dimensionality reduction and clustering of scRNAseq data. In brief, the combined count matrix data was loaded in to R (version 4.0.1) scaled by a size factor of 10,000 and subsequently log transformed. Gene expression cutoffs were at a minimum 200 and a maximum of 6000 genes per cell for each dataset. Cells with greater than 20% mitochondrial genes were filtered. Individual epithelial and stromal libraries were analyzed to create cell type labels based on known marker gene expression. Seurat’s integration was then used to group cell types from disparate patients, integration anchors were identified across all individual patient library samples, as previously described^44^. Specific markers for each cell type that was annotated was determined using the “FindAllMarkers” function. For the epithelial subset analysis, epithelial cells from all patients were subset and integrated together. Epithelial cell states were clustered as described above and clusters were classified using gene scoring according to the previously described cell states^13^, namely for basal, and myoepithelial cell states. Luminal 1-ALDH1A3, Luminal 1-LTF, Luminal 2-MUCL1, Luminal2-AREG. Marker genes for these epithelial subsets can be found in **Supplementary Table 6,** and were used for gene signature scoring across epithelial cells states**. For** single-cell energy (scEnergy) calculation, we employed the R package as recently described^32^. For gene scoring analysis, we analyzed the gene signature transcriptional prevalence in cell types using Seurat’s “AddModuleScore” function. Differential gene expression analysis was performed for each of the cell types, comparing the transcriptome of cells from control and BRCA1^+/mut^ cells using the “FindMarkers” function, using the wilcox rank sum test. To perform statistical tests of gene signature scores between control and BRCA1^+/mut^ patients, the averages of a gene signature score were taken for each individual patient, and groups of means were compared using t-test of control vs BRCA1^+/mut^ patients.

### Ligand-receptor interaction analysis

In order to quantify potential cell-cell paracrine communication in the VMT, we utilized a list of receptor-ligand interactions complied by Skelly et al.^45^ that was generated from Ramilowski et al.^46^ A ligand or receptor is defined as ‘expressed’ if 20% of cells in a particular cell type expressed the ligand/receptor at an averaged level of 0.1. Therefore, a receptor/ligand interaction was considered to be expressed when both the receptor and ligand were expressed in 20% of cells at a level equal or greater than 0.1. To define these networks of interaction, we connected any two cell types where the ligand was expressed in one and the receptor in the other. ‘Enhanced’ receptor-ligand interactions were defined as interactions that were unique within BRCA+/mut or Control cells. To plot networks, we used the chorddiagram function in the R package ‘circilize’. GO Term analysis from receptor-ligand interactions were determined using the gene list enrichment analysis tool ‘Enrichr’^47^ , analyzing unique BRCA or control receptor ligand pairs.

### Human mammosphere growth assay

Human mammary epithelial cells were harvested from cryopreserved organoids that were dissociated into single cells using trypsin, and FACS-isolated based on EPCAM expression. For mammosphere assays, 1x10^4^ FACS-isolated primary human mammary epithelial cells were seeded and suspended in Corning Matrigel Matrix - Growth Factor Reduced (Corning, 354230) and immersed in EpiCult-B Medium (STEMCELL Technologies, 05610) supplemented with 10 ng/mL human recombinant EGF (PeproTech, AF-100-15), 10 ng/mL, 10 ng/mL human recombinant bFGF (PeproTech, 100-18B), 5% FBS (v/v), and 1% Pen Strep (Hyclone, SV30010) (v/v). For cultures treated with exogenous human recombinant NGF (Peprotech 450-01) and human recombinant MMP3 (Peprotech 420-03), 100 ng/mL and 0.5 µg/mL or 1 µg/mL were used in Mammary Epithelial Growth Medium (Lonza, CC-3150), respectively. All cells were grown at 37 °C and at 5% CO2. For quantification of sphere formation, after 5 or 10 days of culture spheres were quantified (in triplicate wells) by imaging the entire matrigel droplet using a BZ-X700 Keyence fluorescent microscope (Keyence Corporation) and manually quantified for number of spheres. Statistics were performed using GraphPad Prism software.

### Epithelial and stromal cell co-culture assay

Fibroblasts/stromal cells were isolated from primary human breast organoid preparations using FACS (EPCAM^low^, CD49^low^,PDPN^+^). Fibroblasts were initially cultured in Fibroblasts Medium (FM) (ScienceCell, 2301) to expand cell numbers. For co-culture experiments, fibroblast cultures up to passage #5 were used. In brief, per well 4,000 primary human mammary epithelial cells were mixed together with 1x10^4^ control, MMP3-overexpression or -knockout fibroblasts and suspended in Corning Matrigel Matrix - Growth Factor Reduced (Corning, 354230) followed by immersion in EpiCult-B Medium for 5-day co-culture experiments. Quantification of mammosphere formation was done as detailed above by imaging. Fibroblasts were identified in culture based on GFP expression and therefore not included in the quantification. Antibodies used for FACS Isolation are listed in **Supplementary Table 11**.

### Lentiviral transduction of primary human stromal cells

Primary mammary fibroblasts were transfected for 48 hours with lentiviral particles with a multiplicity of infection (MOI) of 10 with 10 μg/mL polybrene (Sigma-Aldrich, TR-1003-G).

Lentiviral particles were packaged and purchased from VectorBuilder Inc., Chicago, IL, USA, and contain the following vectors: a GFP expression vector (Cat. No. VB190812- 1255tza), a human MMP3 expression vector (Cat. No. VB170623-1025nbv), a mouse MMP3 expression vector (Cat. No. VB190814-1162wgk), a gRNA expression vector targeting human MMP3 (Cat. No. VB170623-1031qnn), and a Cas9 expression vector (Cat. No. VB170830-1178xap). Transfected cells were isolated by FACS with a BD FACSAriaTM Fusion (Becton Dickinson). For CRISPR/Cas9-mediated MMP3 knockout studies, human primary mammary fibroblasts were first transduced to express GFP and a gRNA targeting human MMP3 and were isolated by FACS using the GFP marker. Subsequently, these cells were expanded and transduced a second time to express mCherry/Cas9 and were isolated by FACS using the mCherry marker. Deletion of MMP3 was confirmed using Western blot as detailed below.

### Human breast morphogenesis assay

Hydrogel branching assays were adapted from a previously described protocol^24, 44^. On ice, Rat Tail Collagen (Millipore, Cat 08-115, Lot# 3026722) is diluted with Lonza Mammary Epithelial Growth Medium (MEGM, Cat# CC-3150) to a concentration of 1.7mg/ml, in NGF treatment 100ng/ml of recombinant NGF (Peprotech 450-01) is diluted in the media. 0.1N NaOH is added to a final pH of 7.2. The extracellular matrix components are added at the final concentrations of 0.5mg/ml of Laminin (ThermoFisher 2301-015), 0.25mg/ml of Hyaluronan (R&D GLR004), and 0.5mg/ml of Fibronectin (ThermoFisher PHE0023). Patient breast tissue that was processed as described in human mammary primary cell isolation, is thawed and washed and loaded into the hydrogel. Hydrogels are plated in 96 well glass bottom dishes (Thermofisher 164588), and then incubated for 1hr at 37C. After hydrogels have solidified, MEGM media is added to the hydrogel, and then incubated at 37C at 5% CO2. Primary branch lengths are measured using ImageJ software. The statistical significance of differences between groups of growth curves was determined by the *Comparing Groups of Growth Curves* permutation test, as described previously.^50^

### Gene expression analysis by quantitative PCR

Cells were sorted by FACS as described above and RNA were extracted by using Quick-RNA Microprep Kit (Zymo Research, R1050) following manufacturer’s instructions. RNA concentration and purity were measured with a Pearl nanospectrophotometer (Implen). Quantitative real-time PCR was conducted using PowerUp SYBR green master mix (Thermo Fisher Scientific, A25742) and primer sequences were found in Harvard primer bank and designed from Integrated DNA Technologies. Gene expression was normalized to the GAPDH housekeeping gene. For relative gene expression 2~negΔΔCt values were used and for statistical analysis ΔCt was used. The statistical significance of differences between groups was determined by unpaired t-test using Prism 6 (GraphPad Software, Inc). Primers are listed in **Supplementary Table 12**.

### *In situ* immunofluorescence analysis

Human mammary tissues were fixed in 4% formaldehyde for 24 hours, dehydrated in increasing concentrations of ethanol, cleared with xylene, and embedded in paraffin. 10 μm tissue sections were prepared using a Leica SM2010 R Sliding Microtome (Leica Biosystems). Slides were heated at 60°C for 20 min, cleared with Histo-Clear (National Diagnostics, Cat. No. HS-200, Atlanta, Georgia, USA) with 2 x 5 min incubations, rehydrated with decreasing concentrations of ethanol, washed in ddH2O and PBS, and subjected to antigen retrieval using a microwave pressure cooker with 10 mM citric acid buffer (0.05% Tween 20, pH 6.0). Tissues were blocked in blocking solution (0.1 % Tween 20 and 10% FBS in PBS) for 20 min, incubated with primary antibodies in blocking solution at 4°C overnight, washed in PBS, incubated with secondary antibodies diluted in PBS for 1 hour, and washed in PBS. Slides were mounted with VECTASHIELD Antifade Mounting Medium with DAPI (Vector Laboratories, H-1200) and micrographs were taken with the BZ-X700 Keyence fluorescent microscope (Keyence Corporation). For quantification of *in situ* data, at least 5 random fields of view per patient sample were imaged and cells were counted manually (blinded). Additional details are provided in the respective figure legends. Antibodies are listed in **Supplementary Table 11**.

### Western blot analyses

Protein samples were subjected to gel electrophoresis, transferred to a PVDF membrane and blocked with a 5% w/v BSA PBST (0.1% Tween-20) solution for 1 hour. Membranes were incubated with primary antibodies overnight at 4°C; MMP3 pAb diluted 1:1000 (Proteintech Group Inc., 17873-1-AP), GAPDH mAb diluted 1:1000 (Cell Signaling Technology, Inc., 2118S). Membranes were washed with PBST (0.1% Tween-20) and incubated with secondary antibodies for 1 hour at room temperature; horseradish peroxidase-conjugated goat anti-rabbit IgG secondary antibody diluted 1:2000 (Thermo Fisher Scientific Inc., G-21234). Membranes were washed with PBST (0.1% Tween-20) and imaged with chemiluminescence reagent (Thermo Fisher Scientific Inc., 34095). Densitometry analyses were performed using ImageJ software.

### Mouse strains

NSG mice were purchased from The Jackson Laboratory (Bar Harbor, Maine, USA). Brca1/p53-deficient mice (*Brca1^f^*^11^*^/f11^p53^f5&6/f5&6^Cre^c^*) were established and genotyped as previously described^5^. All mice were maintained in a pathogen-free facility. All mouse procedures were approved by the University of California, Irvine, Institutional Animal Care and Use Committee.

### Stromal-epithelial co-transplantation for BRCA1-driven cancer initiation *in vivo*

*Brca1^f11/f11^p53^f5&6/f5&6^Cre^c^* mice have a median tumor latency of 6.6 months^5^. Thus, pre- neoplastic primary mammary cells were isolated from all mammary glands (#1 to #5) of 6 month old *Brca1^f11/f11^p53^f5&6/f5&6^Cre^c^* donor mice. Mammary glands were mechanically dissociated, digested in 2 mg/mL collagenase type IV (Sigma-Aldrich, St. Louis, MO, USA) for 1 hour, subjected to differential centrifugation, and digested to single cells with trypsin. Primary human fibroblasts were isolated by FACS (PDPN+) and subjected to lentiviral transduction to express GFP only (+GFP) or both GFP and mouse MMP3 (+MMP3). Fibroblasts were subsequently isolated by FACS based on GFP expression and further expanded *in vitro*. To determine the effect of fibroblast-MMP3 on BRCA1- driven cancer initiation, three cohort of recipient NSG mice (n = 12 each) were transplanted with: 5x10^5^ pre-neoplastic mammary cells; 5x10^5^ pre-neoplastic mammary cells with 5x10^5^+GFP fibroblasts; 5x10^5^ pre-neoplastic mammary cells with 5x10^5^ +MMP3 fibroblasts. Transplantations were done in 100 µL of a 1:1 solution of PBS and growth factor reduced Matrigel (Corning Inc., Corning, NY, USA) into each of the #4 inguinal mammary glands (bilateral injections) in 4-8 week old NSG mice. Tumors were harvested after six weeks and measured with calipers and a scale. The results of Fisher’s Exact test was generated using SAS software. Copyright © 2020 SAS Institute Inc. SAS and all other SAS Institute Inc. product or service names are registered trademarks or trademarks of SAS Institute Inc., Cary, NC, USA. All other statistics were performed with GraphPad Prism software.

### Kaplan-Meier survival analysis

For overall survival analysis, Kaplan-Meier survival curves were generated using microarray data of primary tumors from n=1764 patients in the KM Plotter database^42^. For the overall survival analysis for the pre-caf gene signature , we used marker genes as generated by the ‘FindMarkers’ function in Seurat (**Supplementary Table 3**). For overall survival analysis of preCAF and control gene signatures, a weighted average was calculated with the ‘Use Multiple Genes’ function in KM Plotter. All Kaplan-Meier plots were generated using the ‘auto select best cutoff’ parameter.

### Single-cell western blots

Single-cell western blot assays were performed using the ProteinSimple Milo platform with the standard scWest Kit (ProteinSimple, San Jose, CA, USA) as per manufacturer’s protocol. scWest chips were rehydrated and loaded with cells at a concentration of ∼1 × 10^5^ of cells in 1 mL suspension buffer. Doublet/multiplet capture rate in scWest chip microwells was determined with light microscopy (<2%, established from >1000 microwells). Cells loaded on scWest chips were lysed for 10 seconds and electrophoresis immediately followed at 240 V. Protein was immobilized with UV light for 4 minutes and scWest chips were probed sequentially with primary and secondary antibodies for 1 hour each. Primary antibodies: rabbit anti-KRT 23 (1:20, Sigma, St. Louis, MO, USA); mouse anti-KRT 18 (1:10, Invitrogen, Carlsbad, CA, USA); mouse anti-KRT 14 (1:10, Invitrogen); rabbit anti-KRT 19 (1:10, GeneTex, Irvine, CA, USA). Secondary antibodies: donkey anti-mouse Alexa Fluor 647 (1:10, ThermoFisher Scientific, Carlsbad, CA, USA) and donkey anti-rabbit Alexa Fluor 555 (1:10, ThermoFisher Scientific). Slides were washed, centrifuge-dried, and imaged with the GenePix 4000B Microarray Scanner (Molecular Devices, San Jose, CA, USA). Data was analyzed using Scout Software (ProteinSimple) and ImageJ. Debris, artifacts, and false positive signals were manually excluded during data analyses.

### Mathematical modeling of breast cancer initiation

For our hierarchical model with sequential mutations in oncogenes, We adopted a cancer stem cell model^34^ with sequential mutations of cancer-driver genes to simulate the progress of tumors. The assumptions of the model include 1) within the same genotype, the stem cells self-renewal and give rise to cancer progenitor cells though cell division; 2) cancer progenitor cells differentiate through asymmetric divisions for a limited number of cell cycles; and 3) epithelial stem cells and cancer progenitor cells can switch their genotypes by acquiring mutations in oncogenes, and these driver mutations further increase the cell division rate. Cancer cell populations are considered to be initiated upon the accumulation of driver mutations. To investigate the pro-proliferative effect of MMP3, we estimated the effect of increased proliferation rate of stem and progenitor cells by 2- fold.

For the random mutation model with stochastic fitness shift, we modified a cancer stem cell model^35^ to allow for stochastic changes of individual cell fitness during cell division, induced by both cancer driver and passenger mutations. We assume that the wild-type fitness score is 1, and the advantageous mutations to cell fitness score (i.e. cancer driver mutations) are far less frequent than silent and deleterious mutations to the cell fitness score. We also assume that the stem cell population follows the Moran Process, where cells with high fitness are more likely to proliferate. Similar to the hierarchical model, stromal cues such as NGF and MMP3 enhance the cell proliferation. In this model the populations with larger proportion of high-fitness progenitor cells are more likely to initiate the cancer.

To calculate relative cancer risk ratio, for each patient i, in the jth simulation of random mutation model over the life span, we first calculated high-fitness ratio *p*_*ij*_ as the percentage of progenitor cells with fitness score larger than 1 in the final fitness distribution. The relative risk ratio *R*_*i*_ for patient i is then defined as the likelihood that *p*_*ij*_ is greater than 0.5 in n=20 simulations. We computed the *R*_*i*_ for a population of n=20 patients in each condition with control and 2-fold proliferation rate.

For numerical simulation, we used the R package DIFFpop^34^ to simulate both the hierarchical and random mutation models. In the hierarchical model, the BRCA1^+/mut^ stem cells are treated as the FixPop class with n=10 cells, all other stem cells and progenitor populations are treated as the GrowingPop class, and the differentiated cells are treated as the DiffTriangle class. In the random mutation model, epithelial stem cells are treated as the FixPop class with n=10 cells, cancer progenitor cells the GrowingPop class, and terminally differentiated cells the DiffTriangle class. The stochastic change in fitness induced by mutations is assumed to follow the double exponential.

## Acknowledgements

We thank Devon Lawson and Xing Dai for carefully reading the manuscript. Thank you to Scott Minh-Quoc Nguyen and Nathan Ryan James for their assistance on this project. This study was supported by funds from the NIH/NCI (1R01CA234496; 4R00CA181490 to K.K., and T32CA009054; T32GM008620; F30CA243419 to K.N.), the American Cancer Society (132551-RSG-18-194-01-DDC to K.K.), the NSF (DMS1763272 to Q.N.), The Simons Foundation (594598 to Q.N.), and a grant from Breast Cancer Research Foundation joint with Jayne Kosinas Ted Giovanis Foundation for Health and Policy (to Q.N.). D.M. was supported by the Canadian Institutes of Health Research Postdoctoral Fellowship. Finally, we are grateful to the late Zena Werb for her continuous interest and support of this project.

## Author Contributions

K.K., K.N., and D.M. designed research and supervised research; K.N., D.M., Q.H.N., J.I- R., M.P., G.H., H.A., J.W., M.R., K.R.D., K.B., C.C., A.M., P.S., and D.J. performed research; X.D., J.R., G.X.Y.Z., C.M.N. and Q.N. contributed new reagents and analytic tools; K.N., N.P., P.Z. and Q.H.N. performed computational analyses; K.N., D.M., and K.K. wrote the paper manuscript, and all authors discussed the results and provided comments and feedback.

## COMPETING FINANCIAL INTERESTS

None declared.

## Data Availability

All data will be accessible at Gene Expression Omnibus, including raw .fastq files and quantified data matrices will be deposited in the GEO database (accession code pending).

---

**Extended Data Fig. 1.**
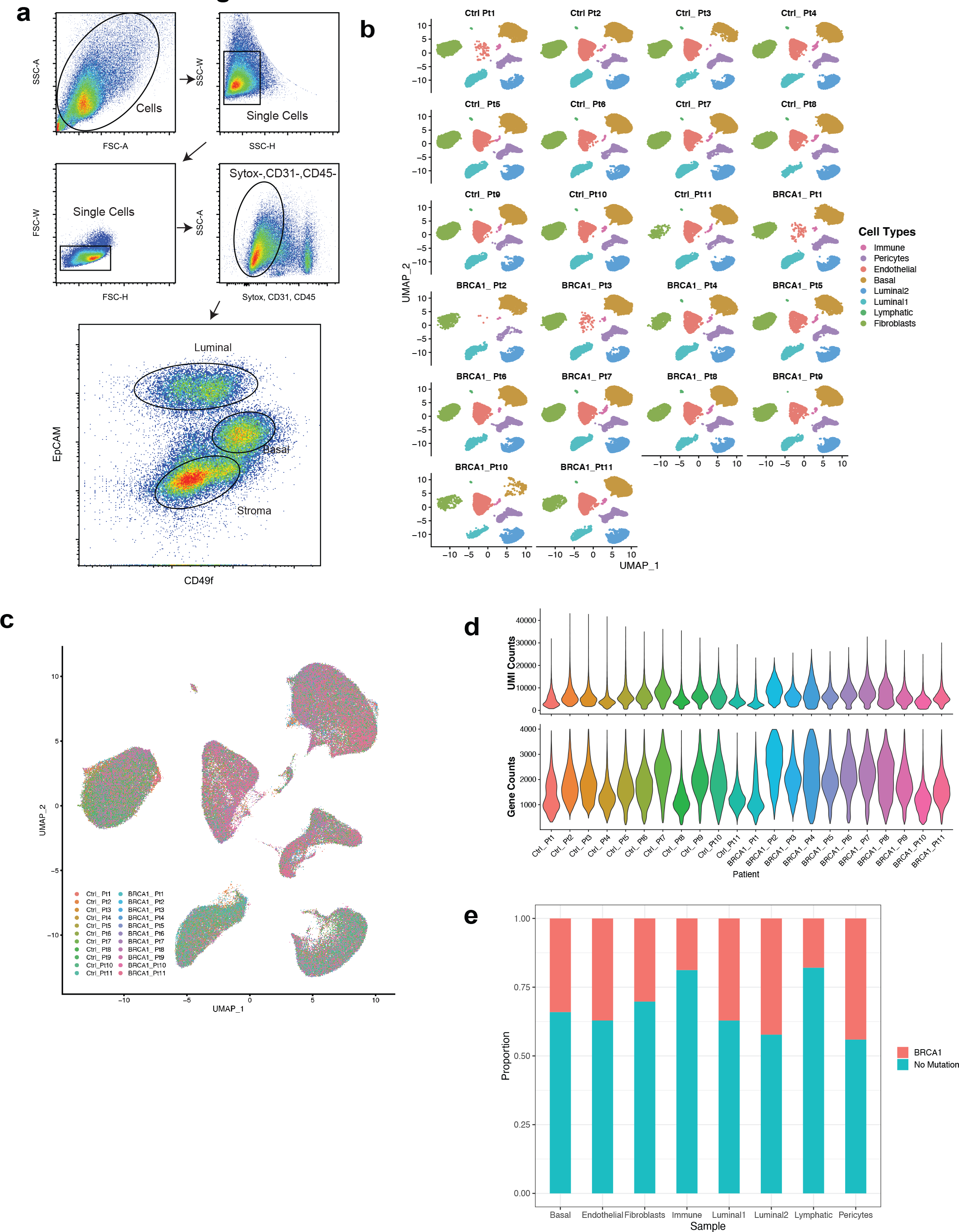
Flow cytometry strategy and quality control metrics for scRNAseq analysis of breast epithelial tissues. a) FACS plots showing gating strategy of mammary epithelial cells in forward and side scatter, singlets gate, dead cell (Sytox+) and lin- (CD31+, CD45+) exclusion gate. FACS plot of the right hand side shows basal (Epcam+, CD49f-high) and luminal epithelial (Epcam-high, CD49f-low) as well as stromal (Epcam-) gating strategy. b) Faceted UMAP projections of n=22 control and BRCA1^+/mut^ patient libraries. Each faceted UMAP projection represents all of the cells from one individual patient. c) UMAP plot labeled by individual. d) Top: Violin plots of UMI counts (the number of individual molecules interrogated/droplet) of each individual patient library. Bottom: Violin plots of number Gene counts (the number of unqiue genes detected/droplet) of each individual patient library e) Bar plots indicating the proportion of each cell type detected in either BRCA1^+/mut^ or control libraries.

**Extended Data Fig. 2.**
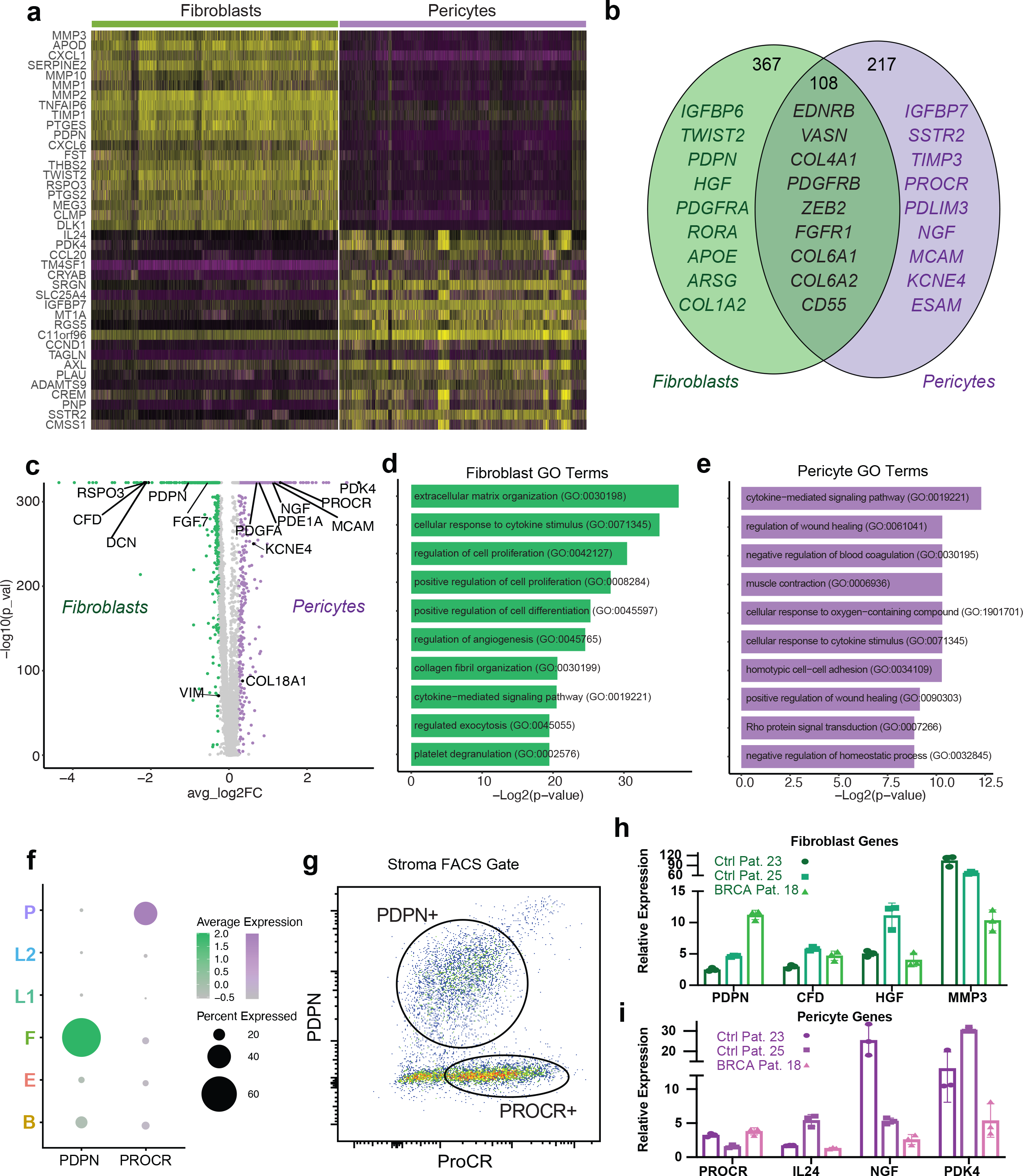
Differential gene expression analysis between fibroblasts and pericytes in the human breast. a) Heatmap showing expression of top 20 marker genes specifically expressed in fibroblasts and pericytes from scRNAseq dataset (rows=genes, columns=cells). Yellow represents a positive z-score, purple represents a negative z-score. b) Venn diagram illustrating the number of genes that are mutually and exclusively expressed in fibroblast and pericyte. Selected marker genes for each category are shown. c) Volcano plot depicting differential gene expression analysis of fibroblasts (green) and pericytes (violet). d) Bar chart showing top 10 GO Terms for fibroblast-specific genes. e) Bar chart showing top 10 GO Terms for pericyte-specific genes. f) Dot plot illustrating selective expression of cell surface markers PDPN and PROCR on fibroblasts and pericytes, respectively. g) FACS plot gated on live cells, singlets, lin-, EpCAM- stromal cells showing distinct populations of PDPN+ and PROCR+ stromal cells. h) Gene expression analysis of FACS-isolated PDPN+ stromal cells by qPCR for selected fibroblast-specific genes. Gene expression normalized to GAPDH and relative expression versus PDPN-PROCR+ pericytes from FACS is shown. i) Gene expression analysis of FACS-isolated PROCR+ stromal cells by qPCR for selected pericyte-specific genes. Gene expression normalized to GAPDH and relative expression versus PROCR^mid^PDPN+ fibroblasts from FACS is shown.

**Extended Data Fig. 3.**
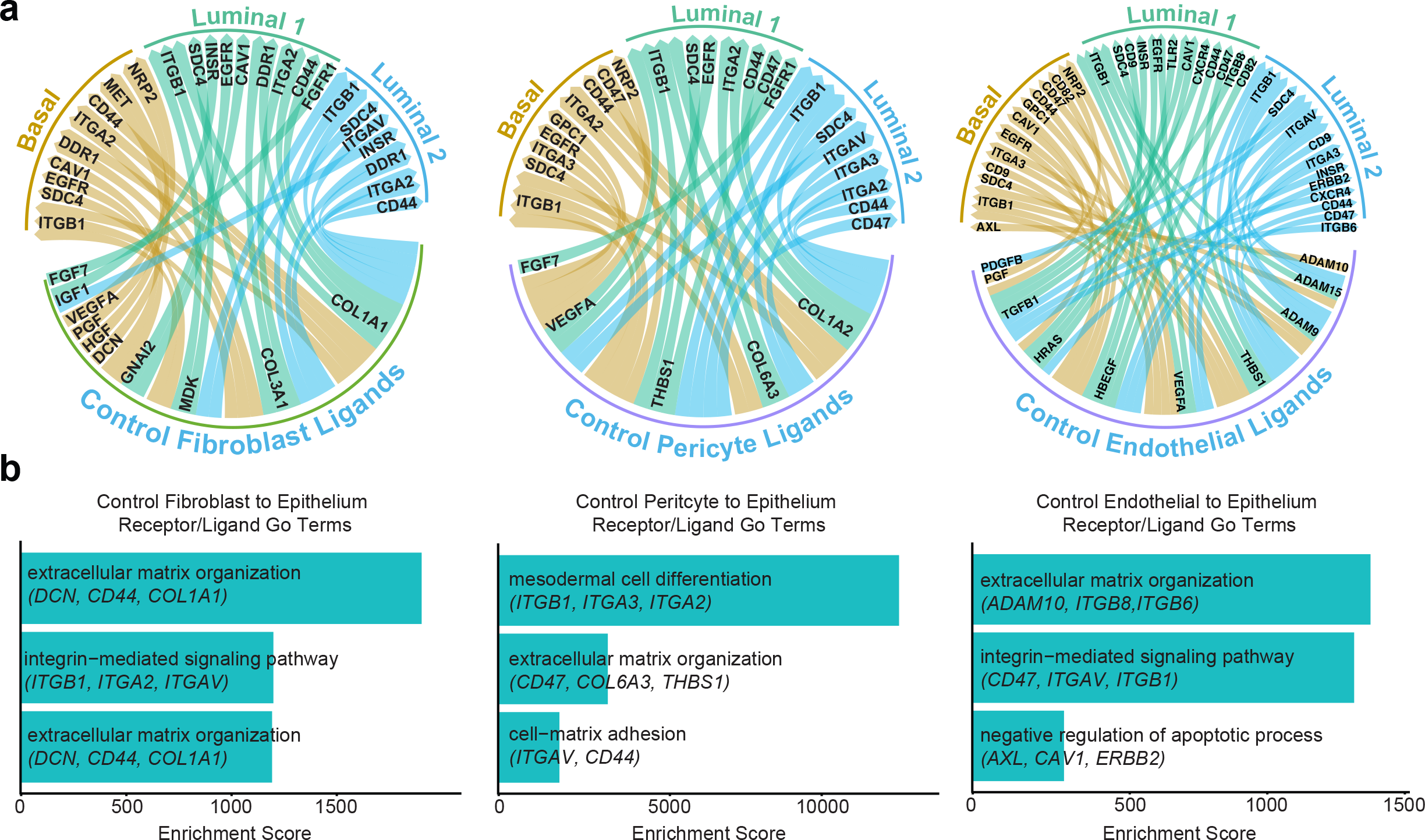
Ligand-receptor interaction analysis analysis of normal tissues. a) Ligand-receptor interactions depicted in circos plots of control fibroblasts, pericyte s and endothelial ligands with interactions to epithelial receptors. b) Receptor-ligand interaction enrichment scores of GO Terms (GO-Biological Processes 2018) of control fibroblasts (left), pericyte (center), and endothelial (right) ligands, and epithelial receptors are shown.

**Extended Data Fig. 4.**
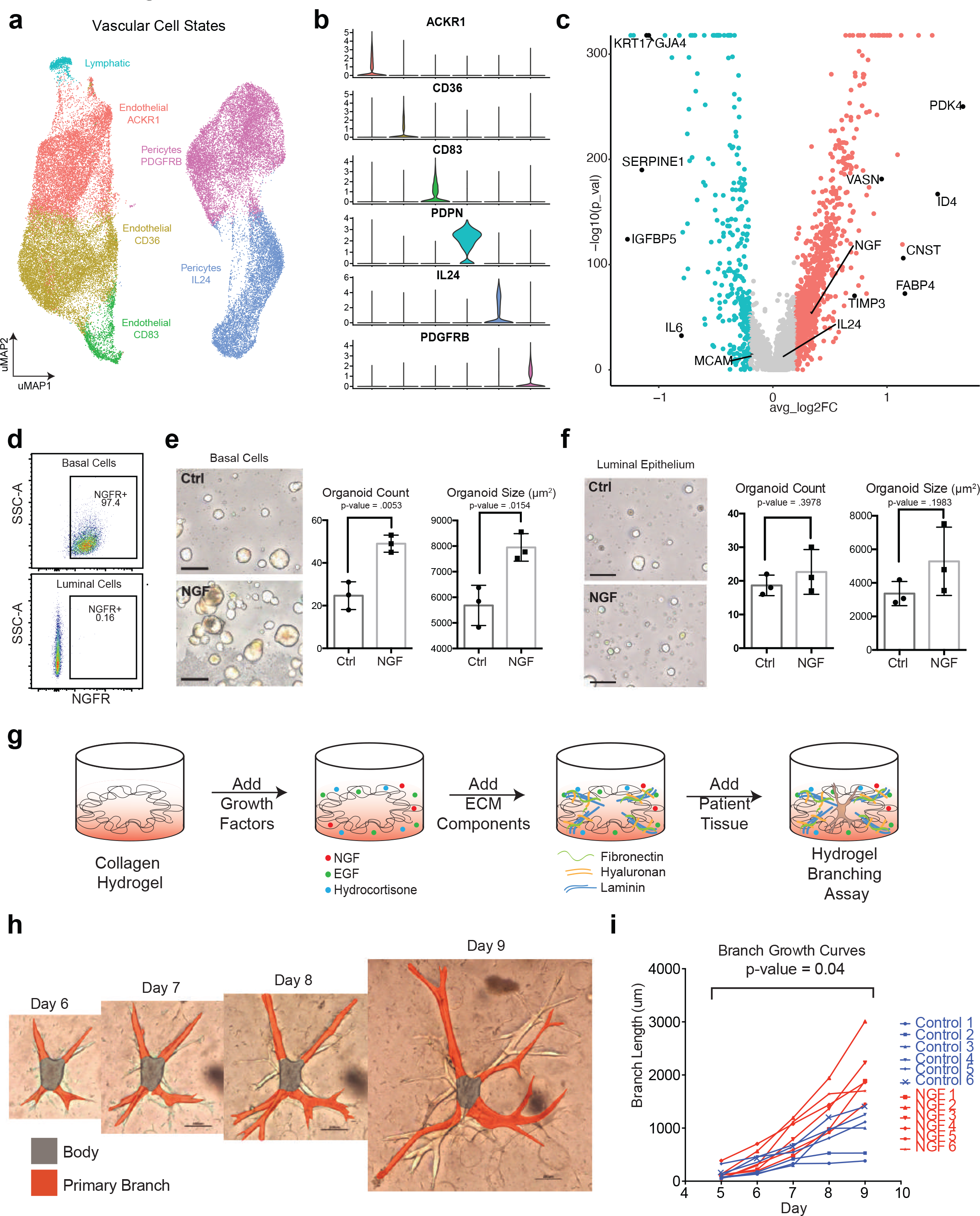
BRCA1+/mut vascular endothelial cells overexpress NGF which increases branching morphogenesis. a) uMAP projection of vascular cell states identifies 3 endothelial cell states, 2 pericyte cell states, and lymphatic cells. b) Violin plots of marker genes with enhanced expression in each vascular cell state cluster. c) Volcano plot showing differentially expressed genes between BRCA1^+/+^ and BRCA1^+/mut^ pericytes. d) FACS plots showing NGFR-expression in basal (top: gated on EpCAM+, CD49f-high) and luminal cells (bottom: gated on EpCAM-high, CD49f- negative). e) Mammosphere assay using FACS-isolated primary human NGFR+ basal epithelial cells grown in the presence of recombinant NGF (100ng/ml) compared to untreated control cells (Ctrl). Representative images are shown on the left, and bar charts showing the number (left) and size (right) of mammospheres in each condition. Error bars depict SD (n=3). *P*-value is determined by unpaired, two-tailed, t-test with equal SD. f) Mammosphere assay using FACS-isolated primary human NGFR- luminal epithelial cells grown in the presence of recombinant NGF (100ng/ml) compared to untreated control cells (Ctrl). Representative images are shown on the left, and bar charts showing the number (left) and size (right) of mammospheres in each condition. Error bars depict SD (n=3). P-value is determined by unpaired, two-tailed, t-test with equal SD. g) Schematic for the generation of hydrogel branching assays h) Time course representative images of a BRCA1^+/mut^ organoid in hydrogel branching assay. i) Branch growth curves of n= 6 control and n= 6 NGF(100ng/ml) treated hydrogel branching assay. P-value = 0.04 was calculated using the CGGC permutation test^43^.

**Extended Data Fig. 5.**
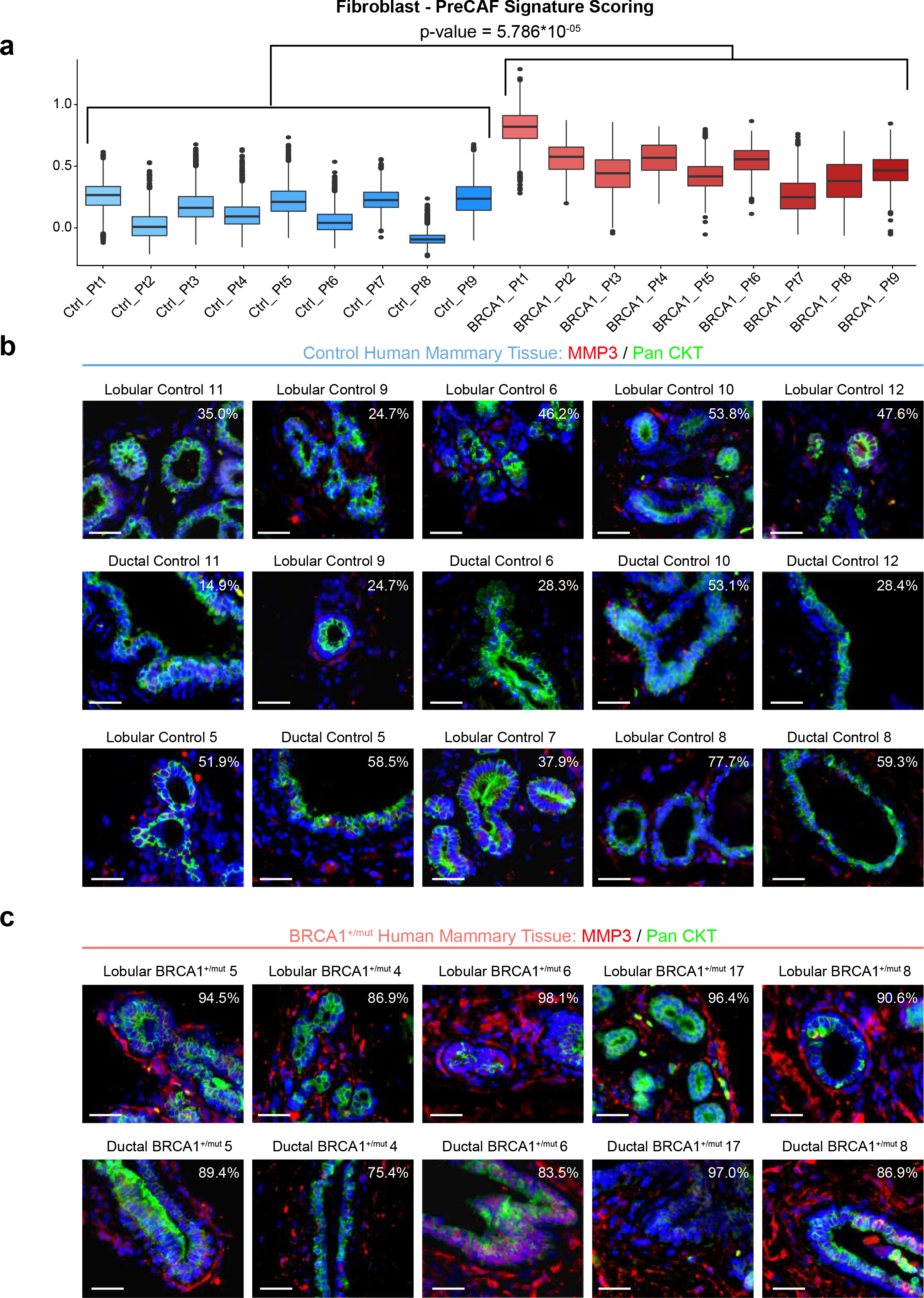
Additional data from *in situ* analysis of MMP3-expressing stromal cells in BRCA1^+/mut^ and control samples. a) Pre-CAF gene signature scoring in fibroblasts from individual patients. Libraries with representation of less than 250 fibroblasts were excluded. *P*-value was determined by a t-test. **a)** Example immunofluorescence microscopy images from ductal and lobular regions in control tissue samples. Values represent the quantified percentage of stromal cells positive for MMP3. Scale bar = 50 µm. **b)** Example immunofluorescence microscopy images from ductal and lobular regions in BRCA1^+/mut^ tissue samples. Values represent the quantified percentage of stromal cells positive for MMP3. Scale bar = 50 µm.

**Extended Data Fig. 6.**
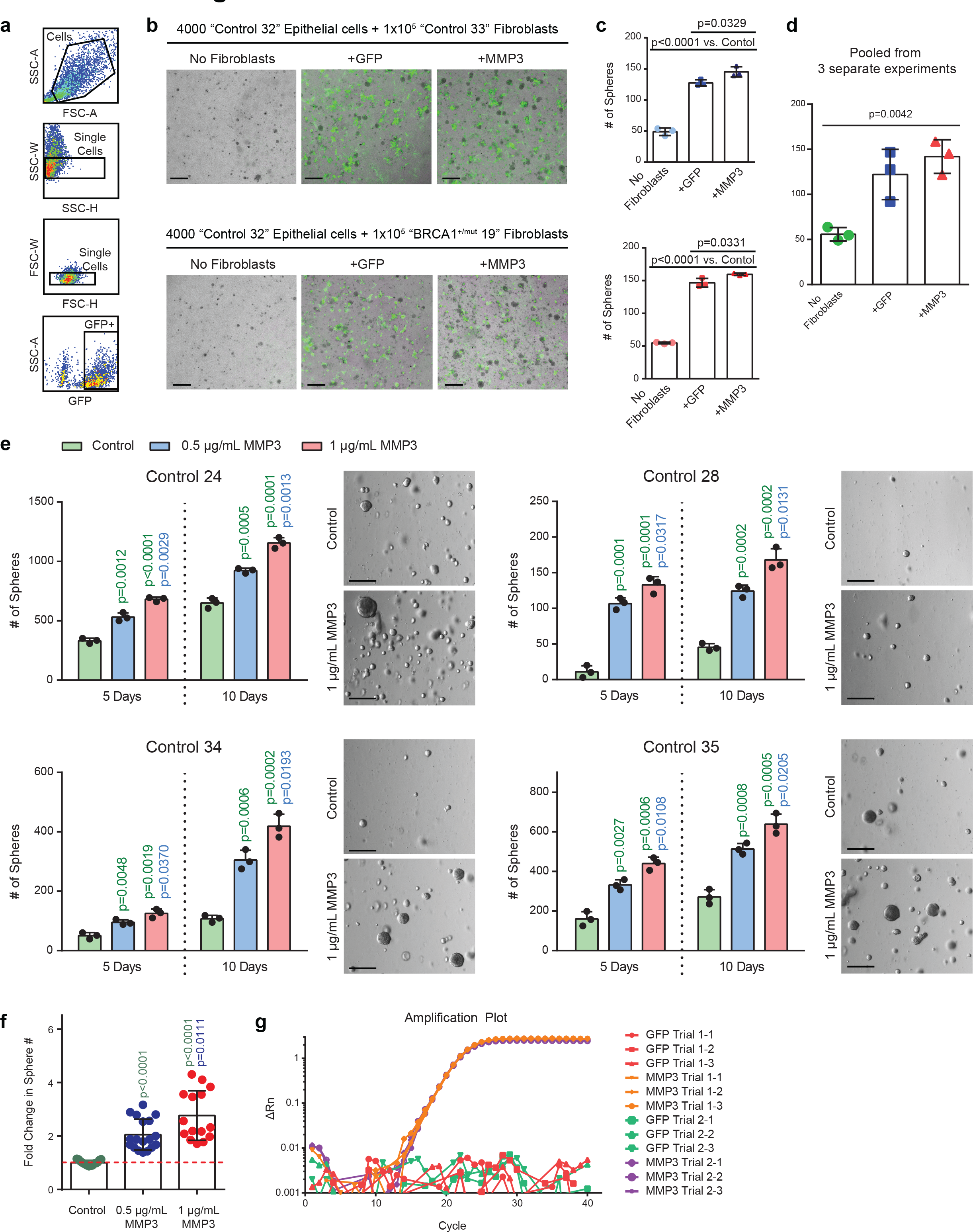
Fibroblast-derived MMP3 promotes epithelial growth *in vitro*. **a)** FACS plots showing gating strategy for isolation of transduced human fibroblasts with GFP fluorescence in forward and side scatter, singlets gate, GFP gate. **b)** Representative images of primary mammary epithelial cells cultured alone (control) or with primary mammary fibroblasts transduced with lentivirus to express GFP only (+GFP) or GFP and MMP3 (+MMP3) in Matrigel for 5 days. Fibroblasts are distinguished from epithelial spheres (GFP negative) with GFP fluorescence. Scale bar = 400 μm. **c)** Quantification of spheres. Values are represented as mean ± SD calculated from nine separate wells per group. P-values were determined by unpaired t-tests. **d)** Mean values of sphere counts pooled from 3 separate experiments from (b) and Fig. 3d. Values are represented as mean ± SD. Statistical significance between all groups was determined with a one-way ANOVA test. **e)** 10x10^4^ FACS-isolated epithelial cells from 4 additional patient samples were seeded in seeded in Matrigel and treated with 0.5 µg/mL or 1 μg/mL MMP3 and spheres were counted after 5 and 10 days. Representative bright field images of mammospheres after 10 days of culture (scale bar= 400 μm). Bar chart values are represented as mean ± SD from triplicates from three separate experiments. *P*- values were determined using unpaired t-tests. **f)** Bar graph depicting the fold change in sphere number compared to control (dotted red line) after 10 days of culture with human recombinant MMP3. Bar graph values are expressed as mean ± SD from 15 independent experiments total (5 different patient samples with 3 separate experiments each). **g)** Primary human breast fibroblasts FACS-isolated from patient sample “Control 27” were transduced to express mouse MMP3 (mMMP3) or GFP only. qPCR analysis was performed on the transduced fibroblasts in two separate trials with three replicates per group. Amplification plot is shown with the difference in the normalized reporter value of the experimental reaction minus the normalized reporter value generated by the instrument (ΔRn) on the y-axis and the cycle number on the x-axis.

**Extended Data Fig. 7.**
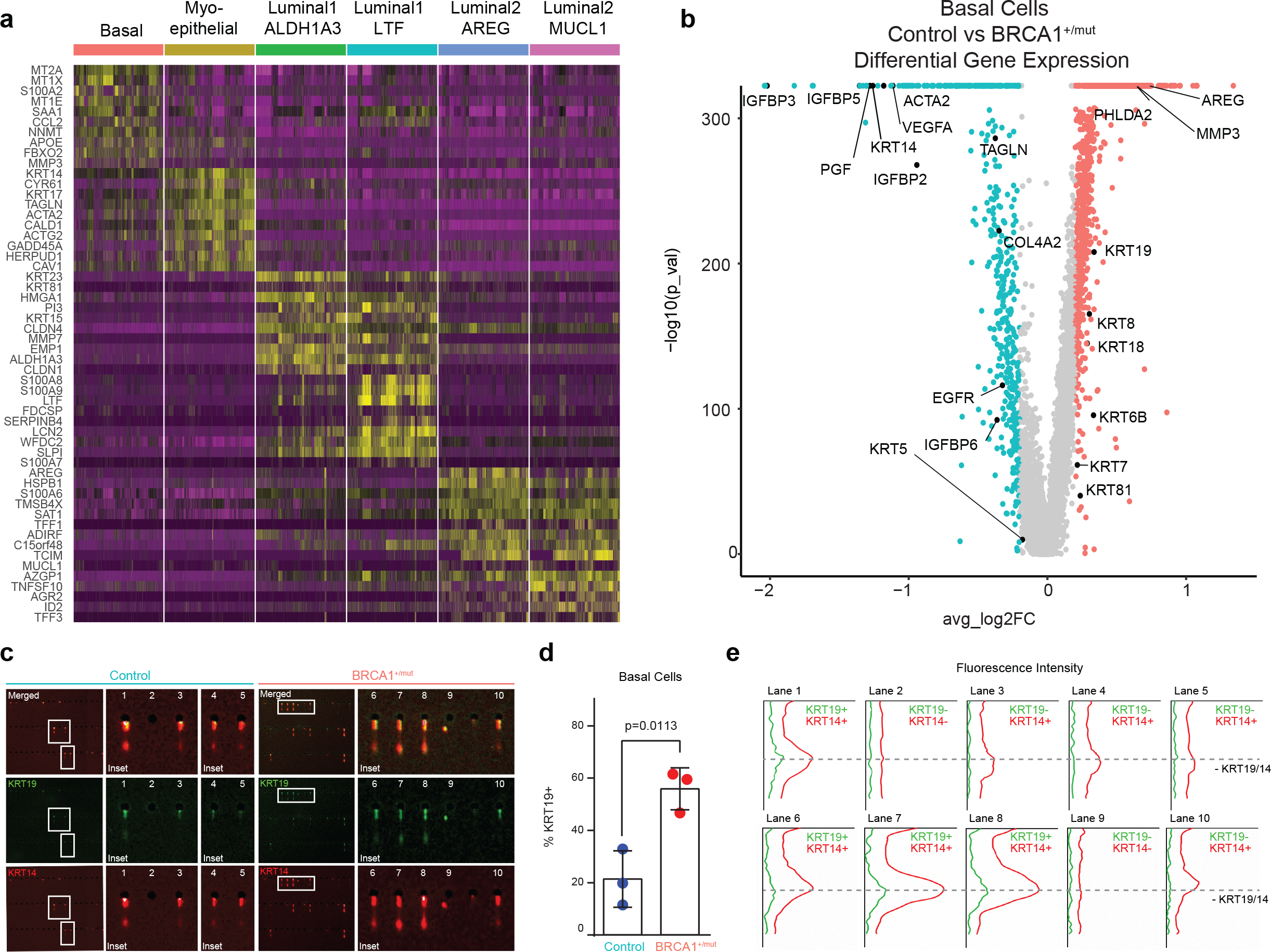
Increased resolution scRNAseq analysis of BRCA1^+/mut^ epithelial cells shows increase of basal epithelial cells with altered differentiation. a) Integrated epithelial clustering analysis of n=11 control and n=11 BRCA1^+/mut^ scRNAseq dataset in uMAP projection showing the main cell types, i.e. the three epithelial cell types basal (B), luminal 1 (L1) and luminal 2 (L2), and the six epithelial cell states. b) Top-10 marker gene heatmap for epithelial cell states. c) Volcano plot showing differentially expressed genes between control and BRCA1^+/mut^ basal epithelial cells. *P* values are determined using the Seurat tobit likelihood-ratio test. d) Single-cell western blot (scWB) analyses for KRT14 and KRT19 were performed on FACS-isolated basal epithelial cells from control and BRCA1^+/mut^ individuals. Representative regions of scWB chips post electrophoresis and antibody probing. e) Quantification of scWBs of all basal cells analyzed. Data is represented at mean ± SD from at least 1000 cells/individual; control n=3, BRCA1^+/mut^ n=3. *P*-value was determined with an unpaired t-test.

**Extended Data Fig. 8.**
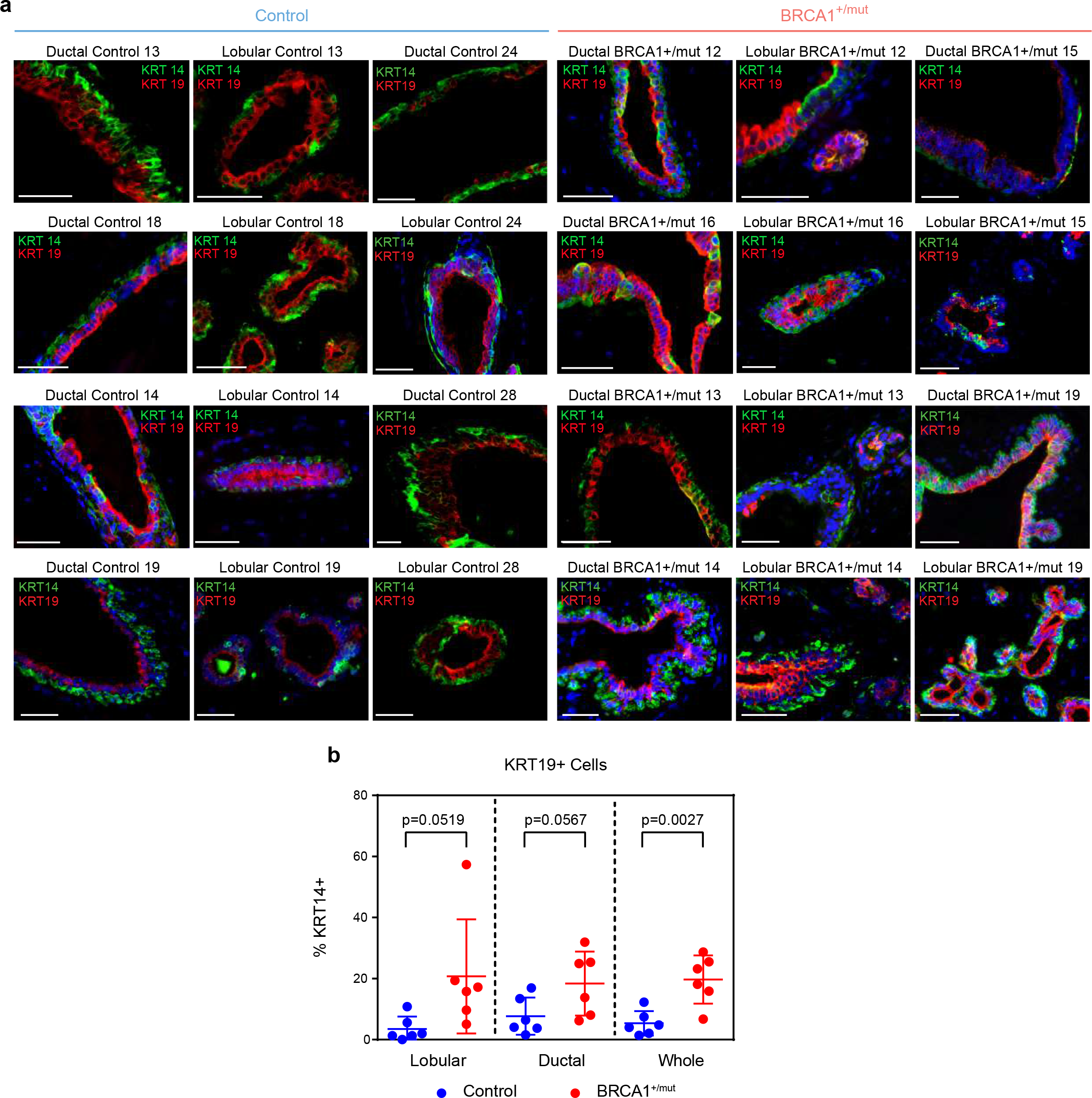
Additional Images Showing BRCA1^+/mut^ have increased number of KRT19+ cells with KRT14 expression. a) Representative immunofluorescence staining for KRT14 and KRT19 in human mammary tissues from control (n=6) and BRCA1^+/mut^ (n=6) individuals. Yellow staining indicates epithelial cells that are double positive. Scale bar= 50 μm. b) Bar graph is a quantitation of the percent of KRT14 and KRT19 double positive cells from lobular and ductal regions of epithelial tissues and whole tissue (lobular + ductal regions). Values are represented as mean ± SD from counts of at least 5 different random fields per tissue. *P*-values were determined by an unpaired t-test.

**Extended Data Fig. 9.**
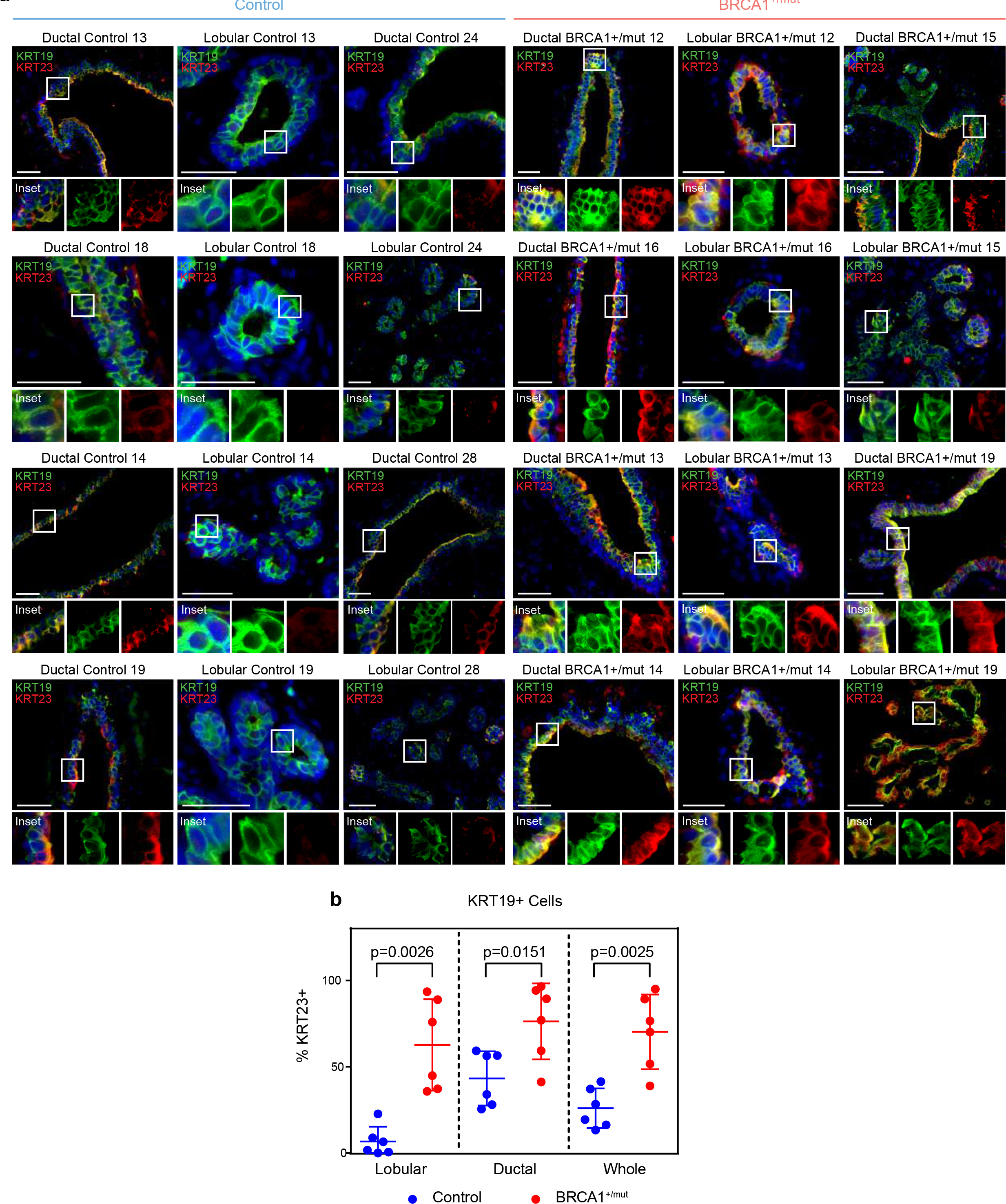
Additional Images Showing BRCA1^+/mut^ have increased number of KRT19+ cells with KRT23 expression. a) Representative immunofluorescence staining for KRT23 and KRT19 in human mammary tissues from control (n=6) and BRCA1^+/mut^ (n=6) individuals. Yellow staining indicates epithelial cells that are double positive. Scale bar= 50 μm. b) Bar graph is a quantitation of the percent of KRT23 and KRT19 double positive cells from lobular and ductal regions of epithelial tissues and whole tissue (lobular + ductal regions). Values are represented as mean ± SD from counts of at least 5 different random fields per tissue. *P*-values were determined by an unpaired t-test.

**Extended Data Figure 10.**
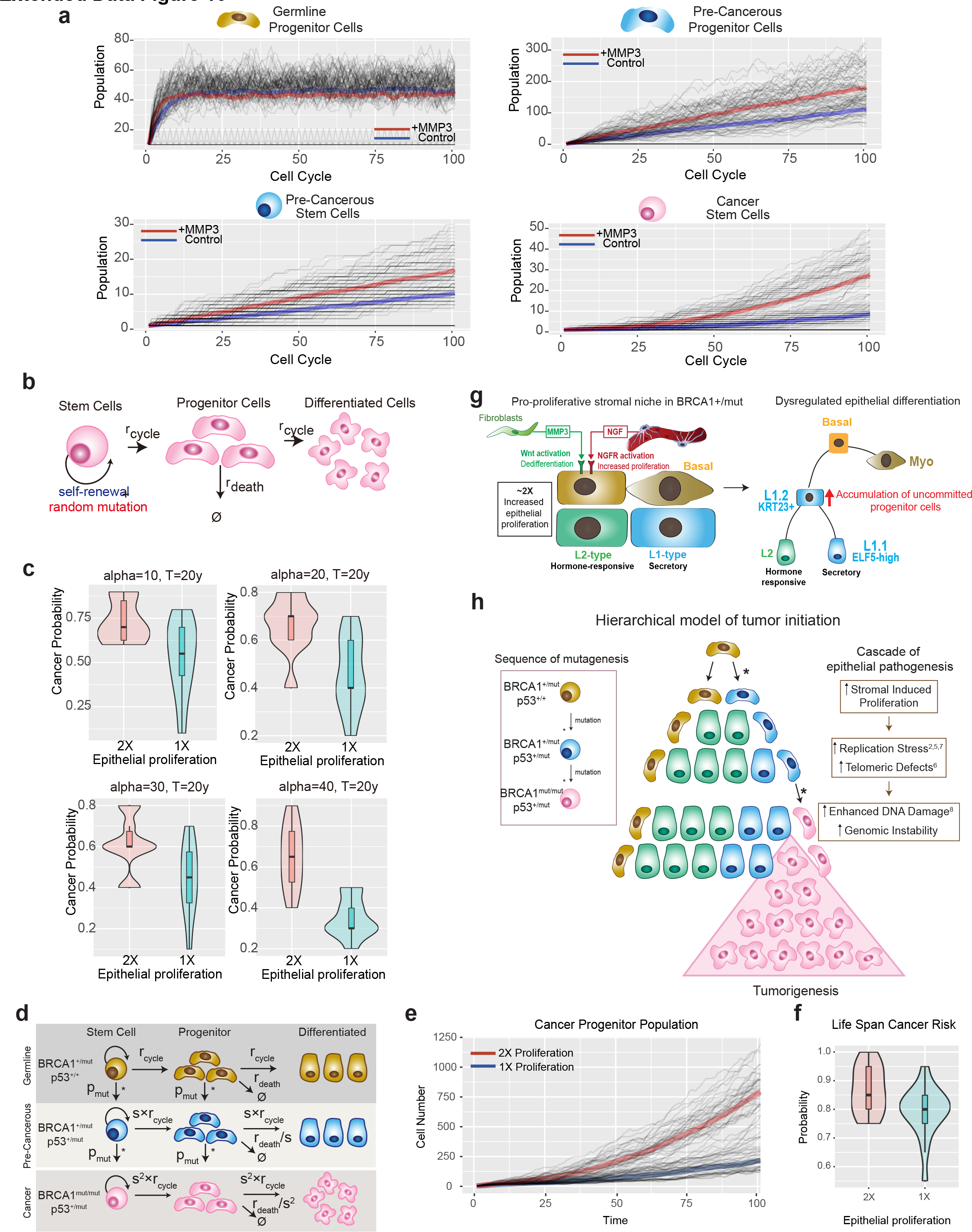
A hierarchical cancer stem cell model predicts increased mutation and cancer initiation risk due to hyperproliferation. a) The comparison of simulated cell population dynamics in 2 fold increased proliferation and control groups for germline progenitor cells (top left), pre-cancerous progenitor cells (top right), pre-cancerous stem cells (bottom left), and cancer stem cells (bottom right). Thick lines: The averaged population dynamics of 2 fold increased proliferation (red) and control group (blue). Gray thin lines: The stochastic simulation trajectories (sample n=50 for each group). b) Schematic model of the assumptions and parameters used to simulate acquisition of random mutations and consequential fitness change in BRCA1+/mut cells. rcycle is the base line cell division rate, rdeath is the cell date rate. Parameters are further defined in **Supplementary Table 10**. c) The robustness of results with respect to parameter *alpha* in the double-exponential distribution of fitness change (**Supplementary Table 10**) in the calculation of risk ratio with or without MMP3. The simulation is performed in the time range of 20 years. d) Schematic model illustrating the assumptions and parameters used to simulate the sequential mutations in oncogenes in BRCA1+/mut cells. rcycle is the base line cell division rate, rdeath is the cell date rate, s = proliferation scale factor, pmut is the probability of acquiring a mutation in a driver oncogene. Parameters are further defined in **Supplementary Table 11**. e) Comparison between cancer progenitor population dynamics as predicted by a hierarchical model^34^ Thick lines: The averaged population dynamics of proliferation in a population with a 2 fold increase in prolifersation and control group (blue). Gray thin lines: The stochastic simulation trajectories (sample n=50 for each group). f) Comparison of the predicted risk ratio of cancer initiation between 2-fold (red) and 1- fold epithelial proliferation rate (blue) in BRCA1 mutation carriers over human life span. The samples are collected from the simulation of N=40 patients in two groups, with the risk ratio of each patient estimated from n=20 simulations of a random mutation model^36^. Violin plots show the distribution of risk ratios over N=20 patients in each group and boxplots indicate median and 25% and 75% quantiles respectively. Wilcoxon Test: *p=0.011. g) Schematic illustrating the concept of a pro-proliferative stromal niche in BRCA1^+/mut^ breast breast tissues. BRCA1^+/mut^ stromal cells overepress pro-proliferative cues including NGF in pericytes and pro-tumorigenic MMP3 in fibroblasts (right side). We propose that these act in concert during the pre-neoplastic phase to promote the expansion of a subset of uncommitted luminal progenitor cells as potential cancer cells of origin (left side). h) Concept illustration of the hierarchical model of cancer initiation. Sequence of mutations are indicated in differently colored cells in box on the left, * represent a mutagenic event. Center schematic summarizes outcome of mathematical modeling results indicating expansion of cancer progenitors and ultimately leading to tumorigenesis. Cascade of epithelial cell-intrinsic events promoting tumorigenesis in BRCA1^+/mut^ is shown on the right. Due to increased stromal cell-induced proliferation and replication stress, BRCA1^+/mut^ breast epithelial stem cells accumulate mutations and become genomically instable, which increases the likelihood of driver mutations, and ultimately tumor initiation.

## Notes

### Competing Interest Statement

The authors have declared no competing interest.

## References

1. Wooster, R. & Weber, B. L. Breast and Ovarian Cancer. N. Engl. J. Med. 348, 2339–2347 (2003).

2. Schlacher, K. et al. Double-Strand Break Repair-Independent Role for BRCA2 in Blocking Stalled Replication Fork Degradation by MRE11. Cell 145, 529–542 (2011).

3. Proia, T. A. et al. Genetic Predisposition Directs Breast Cancer Phenotype by Dictating Progenitor Cell Fate. Cell Stem Cell 8, 149–163 (2011).

4. Lim, E. et al. Aberrant luminal progenitors as the candidate target population for basal tumor development in BRCA1 mutation carriers. Nat. Med. 15, 907–913 (2009).

5. Poole, A. J. et al. Prevention of Brca1-mediated mammary tumorigenesis in mice by a progesterone antagonist. Science 314, 1467–70 (2006).

6. Pathania, S. et al. BRCA1 haploinsufficiency for replication stress suppression in primary cells. Nat. Commun. 5, 5496 (2014).

7. Rosen, E. M. BRCA1 in the DNA damage response and at telomeres. Frontiers in Genetics 4, 85 (2013).

8. Sternlicht, M. D. et al. The stromal proteinase MMP3/stromelysin-1 promotes mammary carcinogenesis. Cell 98, 137–46 (1999).

9. Kessenbrock, K. et al. A Role for Matrix Metalloproteinases in Regulating Mammary Stem Cell Function via the Wnt Signaling Pathway. Cell Stem Cell 13, 300–313 (2013).

10. Inman, J. L., Robertson, C., Mott, J. D. & Bissell, M. J. Mammary gland development: Cell fate specification, stem cells and the microenvironment. Development (Cambridge) 142, 1028–1042 (2015).

11. Speirs, V. et al. Short term primary culture of epithelial cells derived from human breast tumours. Br. J. Cancer 78, 1421–1429 (1998).

12. Macosko, E. Z. et al. Highly Parallel Genome-wide Expression Profiling of Individual Cells Using Nanoliter Droplets. Cell 161, 1202–1214 (2015).

13. Nguyen, Q. H. et al. Profiling human breast epithelial cells using single cell RNA sequencing identifies cell diversity. Nat. Commun. 2018.

14. Crisan, M. et al. A Perivascular Origin for Mesenchymal Stem Cells in Multiple Human Organs. Cell Stem Cell 3, 301–313 (2008).

15. Armulik, A., Genové, G. & Betsholtz, C. Pericytes: Developmental, Physiological, and Pathological Perspectives, Problems, and Promises. Dev. Cell 21, 193–215 (2011).

16. Denu, R. A. et al. Fibroblasts and Mesenchymal Stromal/Stem Cells Are Phenotypically Indistinguishable. Acta Haematol. 136, 85–97 (2016).

17. Sahai, E. et al. A framework for advancing our understanding of cancer-associated fibroblasts. Nat. Rev. Cancer 2020 203 **20**, 174–186 (2020).

18. Agajanian, M., Runa, F. & Kelber, J. A. Identification of a PEAK1/ZEB1 signaling axis during TGFβ/fibronectin-induced EMT in breast cancer. Biochem. Biophys. Res. Commun. 465, 606–612 (2015).

19. Giulianelli, S. et al. FGF2 induces breast cancer growth through ligand-independent activation and recruitment of ERα and PRBΔ4 isoform to *MYC* regulatory sequences. Int. J. Cancer ijc.32252 (2019). doi:10.1002/ijc.32252

20. Matsumoto, K., Umitsu, M., De Silva, D. M., Roy, A. & Bottaro, D. P. Hepatocyte growth factor/MET in cancer progression and biomarker discovery. Cancer Science 108, 296–307 (2017).

21. Descamps, S. et al. Nerve growth factor stimulates proliferation and survival of human breast cancer cells through two distinct signaling pathways. J. Biol. Chem. 276, 17864–70 (2001).

22. Lyu, S., Jiang, C., Xu, R., Huang, Y. & Yan, S. Inhba upregulation correlates with poorer prognosis in patients with esophageal squamous cell carcinoma. Cancer Manag. Res. 10, 1586–1596 (2018).

23. Tirosh, I. et al. Dissecting the multicellular ecosystem of metastatic melanoma by single-cell RNA-seq. Science 352, 189–96 (2016).

24. Macias, H. & Hinck, L. Mammary Gland Development. doi:10.1002/wdev.35

25. Sokol, E. S. et al. Growth of human breast tissues from patient cells in 3D hydrogel scaffolds. Breast Cancer Res. 18, 19 (2016).

26. Shiga, K. et al. Cancer-Associated Fibroblasts: Their Characteristics and Their Roles in Tumor Growth. Cancers (Basel). 7, 2443–2458 (2015).

27. Puram, S. V. et al. Single-Cell Transcriptomic Analysis of Primary and Metastatic Tumor Ecosystems in Head and Neck Cancer. Cell 171, 1611–1624.e24 (2017).

28. Kieffer, Y. et al. Single-Cell Analysis Reveals Fibroblast Clusters Linked to Immunotherapy Resistance in Cancer. Cancer Discov. 10, 1330–1351 (2020).

29. Györffy, B. et al. An online survival analysis tool to rapidly assess the effect of 22,277 genes on breast cancer prognosis using microarray data of 1,809 patients. Breast Cancer Research and Treatment 123, 725–731 (2010).

30. Radisky, D. C. et al. Rac1b and reactive oxygen species mediate MMP-3-induced EMT and genomic instability. Nature 436, 123–127 (2005).

31. Parrinello, S., Coppe, J. P., Krtolica, A. & Campisi, J. Stromal-epithelial interactions in aging and cancer: Senescent fibroblasts alter epithelial cell differentiation. J. Cell Sci. 118, 485–496 (2005).

32. Jin, S., MacLean, A. L., Peng, T. & Nie, Q. scEpath: energy landscape-based inference of transition probabilities and cellular trajectories from single-cell transcriptomic data. Bioinformatics 34, 2077–2086 (2018).

33. Molyneux, G. et al. BRCA1 Basal-like Breast Cancers Originate from Luminal Epithelial Progenitors and Not from Basal Stem Cells. Cell Stem Cell 7, 403–417 (2010).

34. Sedic, M. et al. Haploinsufficiency for BRCA1 leads to cell-type-specific genomic instability and premature senescence. Nat. Commun. 6, 1–14 (2015).

35. Shalabi, S. F. et al. Evidence for accelerated aging in mammary epithelia of women carrying germline BRCA1 or BRCA2 mutations. *Nat*. Aging 2021 19 **1**, 838–849 (2021).

36. Bach, K. et al. Time-resolved single-cell analysis of Brca1 associated mammary tumourigenesis reveals aberrant differentiation of luminal progenitors. Nat. Commun. 2021 121 **12**, 1–11 (2021).

37. Konishi, H. et al. Mutation of a single allele of the cancer susceptibility gene BRCA1 leads to genomic instability in human breast epithelial cells. Proc. Natl. Acad. Sci. U. S. A. 108, 17773–17778 (2011).

38. Ferlic, J., Shi, J., McDonald, T. O. & Michor, F. Genetics and population analysis DIFFpop: a stochastic computational approach to simulate differentiation hierarchies with single cell barcoding. doi:10.1093/bioinformatics/btz074

39. Eyre-Walker, A. & Keightley, P. D. The distribution of fitness effects of new mutations. Nature Reviews Genetics 8, 610–618 (2007).

40. Foo, J., Leder, K. & Michor, F. Stochastic dynamics of cancer initiation. Phys. Biol. 8, 015002 (2011).

41. Pal, B. et al. A single-cell RNA expression atlas of normal , preneoplastic and tumorigenic states in the human breast. 1–23 (2021). doi:10.15252/embj.2020107333

42. Hu, L. et al. Single-Cell RNA Sequencing Reveals the Cellular Origin and Evolution of Breast Cancer in BRCA1 Mutation Carriers. Cancer Res. 81, 2600– 2611 (2021).

43. Coussens, L. M., Fingleton, B. & Matrisian, L. M. Matrix metalloproteinase inhibitors and cancer: Trials and tribulations. Science 295, 2387–2392 (2002).

44. Stuart, T. & Satija, R. Integrative single-cell analysis. Nat. Rev. Genet. 1 (2019). doi:10.1038/s41576-019-0093-7

45. Skelly, D. A. et al. Single-Cell Transcriptional Profiling Reveals Cellular Diversity and Intercommunication in the Mouse Heart. Cell Rep. 22, 600–610 (2018).

46. Ramilowski, J. A. et al. A draft network of ligand–receptor-mediated multicellular signalling in human. Nat. Commun. 6, 7866 (2015).

47. Kuleshov, M. V. et al. Enrichr: a comprehensive gene set enrichment analysis web server 2016 update. Nucleic Acids Res. 44, W90–W97 (2016).

48. Sokol, E. S. et al. Growth of human breast tissues from patient cells in 3D hydrogel scaffolds. Breast Cancer Res. 18, 19 (2016).

49. Miller, D. H., Sokol, E. S. & Gupta, P. B. 3D Primary Culture Model to Study Human Mammary Development. in 139–147 (Humana Press, New York, NY, 2017). doi:10.1007/978-1-4939-7021-6_10

50. CM, E. et al. Leishmaniasis host response loci (lmr1-3) modify disease severity through a Th1/Th2-independent pathway. Genes Immun. 5, 93–100 (2004).

